# Non-apical plateau potentials and persistent firing induced by metabotropic cholinergic modulation in layer 2/3 pyramidal cells in the rat prefrontal cortex

**DOI:** 10.1101/2023.11.02.565356

**Authors:** Nicholas Hagger-Vaughan, Daniel Kolnier, Johan F. Storm

**Author notes:** These authors contributed equally to this work (Nicholas Hagger-Vaughan, Daniel Kolnier).

## Abstract

The prefrontal cortex (PFC) is important for executive functions, including attention, planning, decision-making, and memory, and is proposed by some leading theories to be crucial for consciousness. In particular, the global neuronal workspace theory proposes that PFC layer 2/3 pyramidal cells (L2/3PCs) contribute crucially to the ‘global workspace’, and hence to consciousness, due to their long-range connections to other cortical areas.

Plateau potentials, periods of depolarisation with action potential firing outlasting the stimuli that induced them, have been suggested to help maintain working memory and to contribute to executive functions and consciousness.

We therefore investigated plateau potentials and their mechanisms in PFC layer 2/3 pyramidal neurons. Using whole-cell somatic recordings from L2/3PCs in rat PFC brain slices, we found that the metabotropic cholinergic agonist muscarine reliably induced long-lasting plateau potentials with spiking following a train of evoked action potentials. Similar plateaus were induced by a metabotropic glutamate receptor (mGluR) agonist. Pharmacological tests suggested that these plateaus were dependent on transient receptor potential (TRP) cation channels, both TRPC4 and TRPC5, and required the presence of external calcium (Ca^2+^) and internal Ca^2+^ stores, but not voltage-gated Ca^2+^ channels. Using local Ca^2+^ applications, we found that the responsible Ca^2+^ influx is most likely distributed on the somatic and/or basal dendritic compartments rather than on the (distal) apical dendrite. We used knife cuts to disconnect apical dendrites, sometimes less than 50 µm from the soma, and found that the plateaus did not depend on the distal apical dendrite, since truncated cells generated plateaus with as many spikes as control cells. These results indicate that layer 2/3PCs can generate plateau potentials with sustained spiking independently of distal apical dendrites.

## Introduction

Plateau potentials (PPs) are long-lasting (typically > 100 ms) membrane potential depolarisations that are generated and sustained by intrinsic membrane properties even after the stimulus that triggered them has been terminated (Binder et al., 2009). They can trigger persistent spike firing (PF) or enhance the probability of such firing that can continue for several seconds, sometimes requiring an inhibitory input to switch it off (Egorov et al., 2002). PPs have been observed in a variety of neuron types, and have been suggested to be involved in several functions, including working memory, neural plasticity, associative learning, and conscious perception (Aru et al., 2020; Fransén et al., 2006; Gambino et al., 2014; Hasselmo & Stern, 2006)

There is considerable evidence for PPs or other intrinsically generated sustained depolarisations of dendritic origin, for example in the apical dendrites of neocortical pyramidal cells, generated by dendritic voltage-gated Ca^2+^ channels (VGCCs) or N-methyl-d-aspartate receptors (NMDARs), and resulting in somatic sustained spiking or bursts (Gidon et al., 2020; M. E. Larkum et al., 1999). PPs have been suggested as a mechanism for maintaining working memory (Egorov et al., 2002; Fransén et al., 2006; Hasselmo & Stern, 2006), and shown to be important for induction of long-term potentiation (LTP) in apical dendritic synapses in vivo (Gambino et al., 2014).

Apical dendritic calcium spikes and PPs have also been posited as a mechanism for integration of different information streams, which is needed for conscious sensory perception according to the dendritic integration theory of consciousness (Aru et al., 2020; Suzuki & Larkum, 2020). Thus, in rodent layer 5 (L5) pyramidal cells, simultaneous synaptic input to the perisomatic region and the distal apical dendrite can cause a dendritic Ca^2+^ spike/PP that triggers an output of axonal/somatic bursts of Na^+^ action potentials (M. E. Larkum et al., 1999). The ‘apical amplification’ caused by coincident synaptic inputs from sensory (i.e. thalamic, perisomatic targeting) and contextual (cortico-cortical, apical targeting) information streams provide, per the theory, a key mechanism for conscious perception (Aru et al., 2020; Phillips et al., 2016; N. Takahashi et al., 2016).

PPs and PF can often be induced or enhanced by neuromodulation, e.g. by acetylcholine (ACh) via muscarinic receptors (mAChRs; (Fraser & MacVicar, 1996; Rahman & Berger, 2011), dopamine (Baufreton et al., 2003; Hernández-López et al., 1997), serotonin (Perrier & Hounsgaard, 2003; D. Wang et al., 2014), or via metabotropic glutamate receptors (mGluRs) (Yoshida et al., 2008). Neuromodulator-induced PPs and PF have been found in a variety of excitatory neuron types, in several brain regions, including the hippocampus (Hagger-Vaughan & Storm, 2019; Kuzmiski & MacVicar, 2001) and neocortex (Heng et al., 2011).

The prefrontal cortex (PFC) is considered to be of key importance for executive functions, including planning (Tanji & Hoshi, 2001), decision-making (Hunt, 2021), and outcome prediction (Del Arco et al., 2017). The PFC is also posited as being essential for consciousness in several leading theories, including the global neuronal workspace theory (Mashour et al., 2020) and the higher-order theory consciousness (Brown et al., 2019).

Layer 2/3 pyramidal cells (L2/3PCs) in the PFC have been postulated to have a key role in consciousness, in particular in the GNWT, where the long-range cortico-cortical connections formed by L2/3PCs provide the architecture required for the “global work-space" and its “ignition” and reverberant network activity, which is considered essential for conscious access (Brown et al., 2019; Mashour et al., 2020). Whilst persistent network activity is likely dependent on recurrent synaptic connections (X. J. Wang, 2001), intrinsic cellular properties promoting enhanced and sustained firing are likely often necessary to support this network activity (Jochems & Yoshida, 2015). Thus, in a network model of the hippocampal CA3 network, persistent network activity was supported by a combination of synaptic changes and intracellular PP mechanism (Jochems & Yoshida, 2015) induced by mAChR activation (Jochems & Yoshida, 2013).

This suggests that networks can be enabled to support persistent, recurrent activity, which may underpin conscious processing and memory, by intrinsic cellular properties induced by neuromodulators. This is of particular interest in the case of ACh, which is released in the cortex at high levels during wakefulness and dreaming (REM) sleep when there are different forms of conscious experience, in contrast to the unconscious state of dreamless sleep, when the ACh release is far lower (Lee et al., 2005; Watson et al., 2010).

In this study, we described and characterised ACh-induced plateau potentials (PPs) in PFC L2/3 pyramidal cells, and investigated the underlying mechanisms, including which ion channels generate the PPs and which subcellular compartments are needed for their generation.

## Methods

### Animals

All experimental procedures were approved by the responsible veterinarian of the Institute, in accordance with the statute regulating animal experimentation given by the Norwegian Ministry of Agriculture, 1996.) For use in optogenetics experiments (see below, and **Suppl. figure 5**), transgenic Cre reporter allele mice under th e control of the cholinesterase gene (Chat-IRES-Cre; Stock No. 006410) were cross-bred with Ai32 mice (B6.Cg-Gt(ROSA)26Sortm32(CAG-COP4*H134R/EYFP)Hze/J; Stock No. 024109), thus generating experimental animals expressing yellow fluorescent protein (YFP) and channelrhodopsin in cholinergic cells. All transgenic animals were obtained from The Jackson Laboratory.

### Brain slice preparation

Young male Wistar rats (P21-P28) or adult (transgenic) mice of either sex, were briefly anaesthetized with suprane before decapitation and removal of the brain into ice-cold cutting solution containing (in mM): NaCl 87, KCl 1.25, KH_2_PO_4_ 1.25, NaHCO_3_ 25, glucose 16, sucrose 75, MgCl_2_ 7.0, CaCl_2_ 0.5. Coronal slices of 350-400 µm thickness were made in ice-cold cutting solution using a VT1200 vibratome (Leica Microsystems, Wetzlar, Germany). The slices were transferred to a submerged holding chamber containing the artificial cerebrospinal fluid (aCSF) used for recording (described below) at 35 °C for 30 minutes before being moved to room temperature (20-24 °C). For experiments with cells with truncated apical dendrites (**Figures 6 & Suppl. Figure 5**), the slice was cut parallel to the pia approximately at the level of the layer 1/layer 2 border, using a scalpel blade immediately prior to recording.

### Electrophysiology

For most recordings slices were submerged in an aCSF containing (mM): NaCl 125, KCl 2.25, KH_2_PO_4_ 1.25, NaHCO_3_ 25, glucose 16, MgCl_2_ 1.5, CaCl_2_ 2.0, DNQX 10 µM, SR 95531 (gabazine) 5 µM, DL-2-amino-5-phosphonovalerate (AP5) 50 µM. For experiments using reduced-Ca^2+^ aCSF, the composition was as above except for the following changes (in mM): MgCl_2_ 4.0, CaCl_2_ 0.1, EGTA 0.2 (**Figures 4 & 5**). Cells were viewed using IR-DIC optics on a BX51WI microscope (Olympus, Tokyo, Japan). Whole-cell, somatic patch-clamp recordings were obtained from L2/3PCs in the paralimbic or infralimbic subregions of the prefrontal cortex. Patch pipettes were pulled from borosilicate glass using a Narashige PC-10 vertical puller (Narashige, Japan). Pipettes had tip resistances of 4-6 MΩ and were filled with a solution containing (in mM): K-methanesulphonate 120, HEPES 10, KCl 20, MgATP 4, NaGTP 0.4, Na_2_phosphocreatine 5, EGTA 1.0. The pH was adjusted to 7.3 and the osmolarity was adjusted to 290 mOsmol/l. For some experiments, Alexa 488 10 µM, Alexa 555 15 µM, or biocytin 0.3% was added to the pipette solution for tracing the cell morphology. The liquid junction potential was 9.4 mV and was not corrected for. Current-clamp recordings were performed using a Multiclamp 700A amplifier (Molecular Devices, Sunnyvale, CA, USA) or a Dagan BVC-700A (Dagan Corp, Dagan, Minneapolis, MN, USA). Local chemical application was achieved using a Picospritzer 2 (Parker, Hollis, NH, USA). Cells were filled with Alexa 555 for up to 30 minutes before fluorescence was observed using a mercury lamp and an Andor Zyla 4.2 sCMOS camera (Oxford Instruments, Abingdon, UK). This was then used to guide the pipette for drug application to the appropriate part of the cell. Same-sized pipettes were used for focal application as for whole-cell patch-clamp. Pipettes used for focal application contained modified aCSF, with the inclusion of (in µM): muscarine 10, Alexa 555 10, DNQX 10, gabazine 5, and AP5 50. For experiments using reduced-Ca^2+^, the following changes were made: 5 mM Ca^2+^, 0.5 mM Mg^2+^. In experiments with focal application of FFA, 200 µM FFA (in DMSO, final concentration in extracellular aCSF was >0.1%) was added. In control experiments, DMSO was used instead of FFA. A 60-s long pressure pulse of 1-10 psi, with the pipette 10-40 µm from the soma, or near the first bifurcation of the apical dendrite, was used, starting 30 seconds before the induced spike-train. For experiments with focal application of FFA and the corresponding control experiments, a 120-s long pulse was used instead, starting 90 seconds before the induced spike-train.

### Optogenetics

Activation of cholinergic afferents occurred by a train of 5 or 10 blue light (λ = 460-480) pulses, 10 ms long, at 25 Hz, from a mercury lamp (Olympus, Tokyo, Japan) or a xenon lamp (Lambda XL, Sutter, Novato, CA, USA) (**Suppl. Figure 5**). Illumination was centred on the soma using either a 40x (NA 0.8) or a 60x (NA 0.9) immersion lens. In all experiments with photoactivation of cholinergic afferents, 10 µM gabazine and 10 µM physostigmine were included in the aCSF.

### Data acquisition and analysis

Data were acquired using pClamp 10.7 software, and the data were digitised using a Digidata 1440A (Molecular Devices). Analysis was performed in Clampfit 10.7 (Molecular Devices), and plotted in Origin 9.7 (OriginLab, Northampton, USA).

### Histology

Cells intended for staining and reconstruction were filled with biocytin during electrophysiological recording and fixed in 4% paraformaldehyde in phosphate-buffered saline overnight at 4 °C, then transferred into phosphate-buffered saline at 4 °C for up to two weeks. The avidin-biotin-peroxidase method was used to visualise the cells, using 3,3′-diaminobenzidine as a chromogen (ABC kit from Vector Laboratories, Burlingame, CA, USA) and morphological reconstruction was performed using the Neurolucida system (MicroBrightField, Colchester, VT, USA).

### Chemicals

DNQX, gabazine, DL-AP5, ML 204, AC 1903, nifedipine, muscarine, tACPD and physostigmine were acquired from Tocris. All chemicals were bath applied at a superfusion rate of ∼2 ml min^−1^, except where described otherwise. Note that in experiments with bath-applications of drugs/chemicals or modified aCSF (e.g. in **Figures 2-4**), we normally compared the changes seen after the bath-application (i.e. after a time giving apparently full effect; typically after a few minutes) with changes seen in time-matched control recordings (usually in different cells), when no chemical was applied and the aCSF was not changed. This is a more informative comparison than simply comparing parameters before and after each application, because several parameters can change spontaneously over time, even when the aCSF was not changed.

### Statistics

Statistical analysis was performed with GraphPad Prism, version 9. Grouped data are expressed as mean ± SEM, and the sample size of cells (*n*). Non-parametric tests were used to determine statistical significance as sample sizes were too small to accurately determine whether they were normally distributed. Paired data were checked for significance using the Wilcoxon Signed-Rank (WSR) test, whilst the Mann-Whitney U (MWU) test was used for unpaired data. Summary graph plots include all data points as well as mean lines and 95% confidence intervals (CI) represented by whiskers. Differences between groups were considered significant at *p* < 0.05 and are denoted in graphs by an asterisk. Group differences with *p* > 0.05 were considered non-significant and are denoted in graphs by “ns”.

## Results

### mAChR-dependent plateau potentials in PFC L2/3PCs

We obtained stable whole-cell recordings from 102 layer 2/3 pyramidal cells from the prelimbic and infralimbic regions of the prefrontal cortex in acute brain slices from rats and mice (**Figure 1A-B**). To elicit spike trains, we used depolarising somatic injections of 7 brief (2 ms) depolarising current (1-1.5 nA) pulses at 70 Hz, each triggering a single action potential (hereafter referred to as the spike-train protocol).

**Figure 1.**
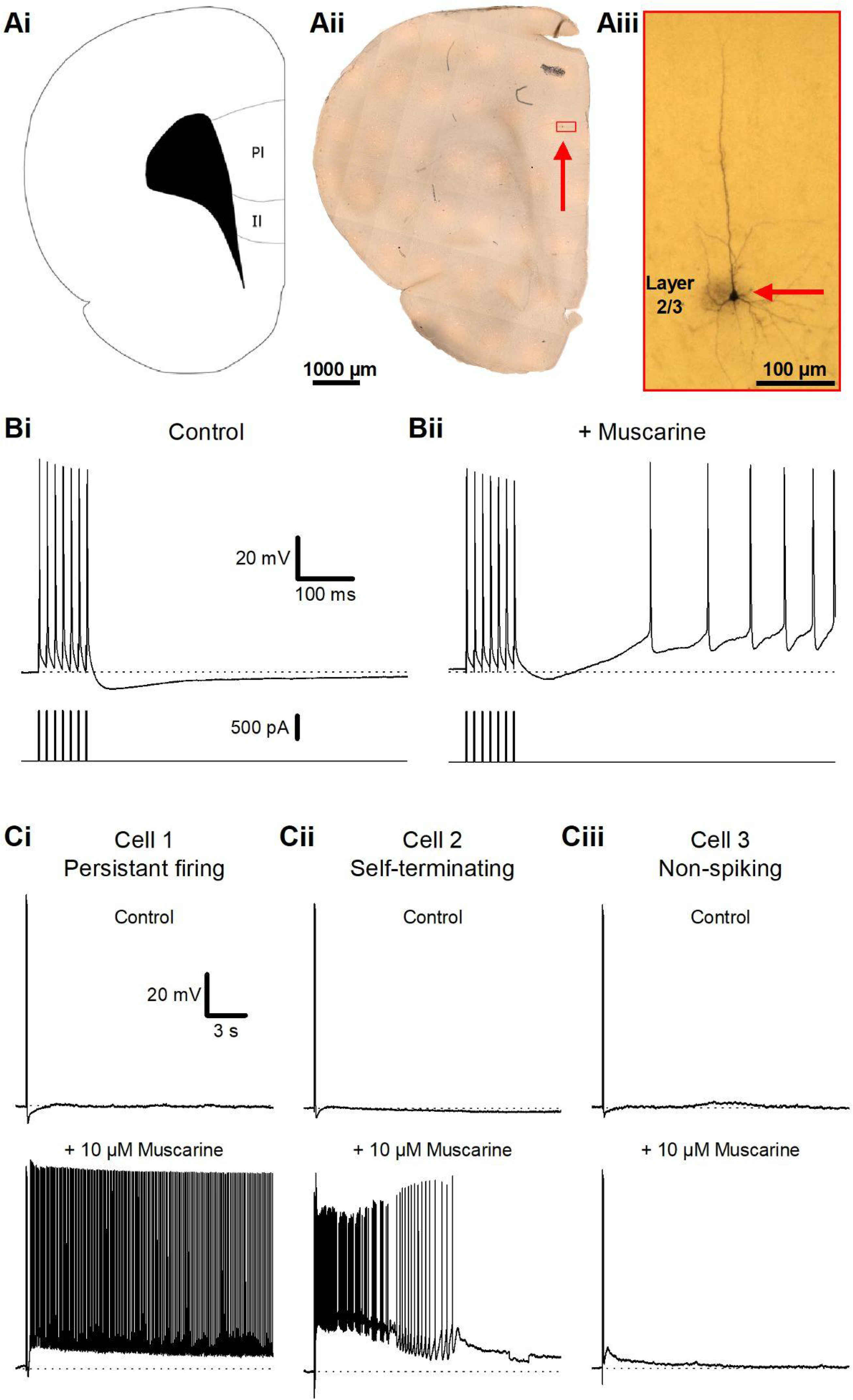
Muscarinic plateau potentials in L2/3PCs of the PFC. **A** - Visual overview over an example recorded cell, with a schematic of a coronal brain slice (i) showing the PFC regions where cells were recorded from (Paxinos & Watson, 2006), a fixed rat PFC slice (ii), shown at low magnification, with a red box marking the location of the biocytin stained L2/3PC seen in (iii). **B** - Somatic whole-cell recording traces from a L2/3PC, showing membrane potential responses to a train of 7 action potentials at 70 Hz (top traces), evoked by injected brief depolarising current pulses (bottom traces), under control conditions (i), and after the bath-application of 10 µM muscarine (ii). **C -** Example traces of different membrane potential responses in three different cells (i-iii) after a train of current pulses during control condition (top traces) and after the bath application of 10 µM muscarine (bottom traces).

In control conditions (normal aCSF) each spike train was followed by a slow after-hyperpolarisation (AHP; **Figure 1Bi**), lasting several hundred milliseconds, but during bath-application of 10 µM muscarine, the spike train elicited a range of depolarising changes in the membrane potential, which varied between cells (**Figure 1C**): from sustained PPs, eliciting spike firing for several seconds (**Figure 1 Ci**) to briefer depolarisations with (**Figure 1 Cii**) or without (**Figure 1 Ciii**) spiking. Here, both long-lasting PPs and briefer ADPs will be referred to as PPs, since they seem to represent different degrees of the same phenomenon, and share key properties and mechanisms (see below).

We tested how PPs depend on the concentration of muscarine (**Figure 2**). The lowest muscarine dose (1 µM) abolished the slow AHP but elicited only a small post-burst depolarisation (PBD; **Figure 2 Ai**) (control: -0,10 ± 0.21 mV; 1 µM muscarine: 0.90 ± 0.25 mV, *n* = 10, *p* = 0.02), which did not trigger any spikes in the ten cells tested. (Note that the values for changes in membrane potential (PDB) were measured 5 minutes after the onset of muscarine application, and are always compared to time-matched values for control recordings, to compensate for any time-dependent changes. The PDB values may thus occasionally be negative.) Tripling the muscarine concentration (3 µM) induced a stronger PBD a (control: -0.56 ± 0.29 mV; 3 µM muscarine: 9.9 ± 2.2 mV, *n* = 9, *p* < 0.01), which in most cells triggered spiking (control: 0 ± 0 spikes; 3 µM muscarine: 69 ± 19 spikes, *n* = 9, *p* = 0.03). Increasing the muscarine concentration to 10 µM gave a still larger PBD (control: -0.36 ± 0.12 mV; 10 µM muscarine: 18 ± 2.0 mV, *n* = 10, *p* < 0.01) and more spiking (control: 0 ± 0 spikes; 10 µM muscarine: 71 ± 11 spikes, *n* = 10, *p* < 0.01) (**Figure 2B & C**). The values compared in **Figure 2C** were measured after 5 minutes of drug application (or no drug, for the controls). The different muscarine concentrations (1, 3, and 10 µM) were tested in different cell groups, not added cumulatively in the same cells.

**Figure 2.**
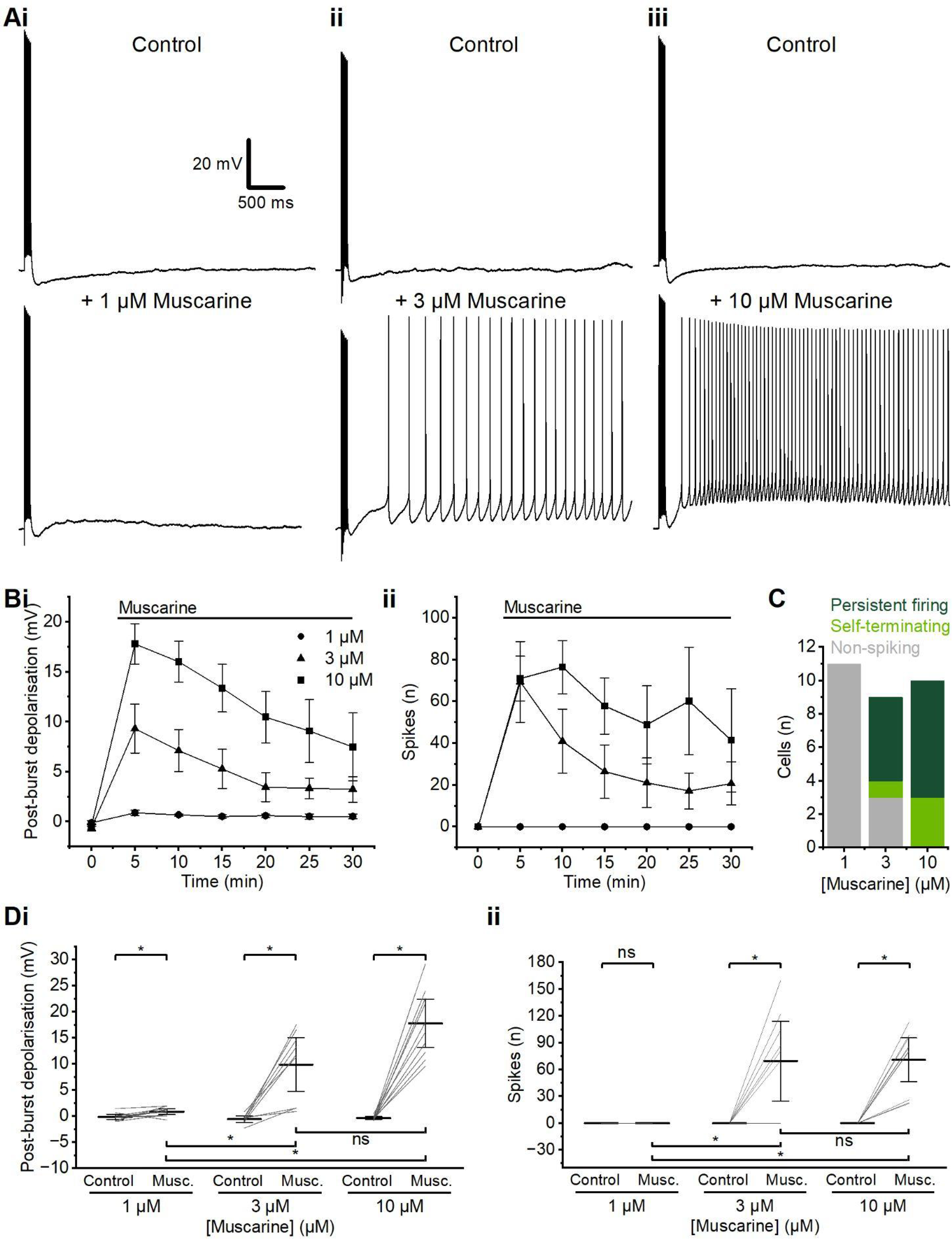
Plateau potentials depended on the concentration of muscarine. **A** - Example traces showing characteristic responses following an action potential train during the application of 1 µM (i), 3 µM (ii), or 10 µM muscarine (iii). **B** - Time course comparison of PBD (i) and spikes (ii) at different muscarine concentrations. **C** - Summary graph of the incidence of persistent firing, self-terminating and non-spiking evoked by different muscarine concentrations. **D** - Summary graphs for the PBD (i) and spikes (ii) evoked by different muscarine concentrations. (i) Data shows a significant increase in PBD at all concentrations tested of muscarine compared to control conditions. (WSR test, 1 µM: *n* = 10; 3 µM: *n* = 9; 10 µM: *n* = 10). Comparison of PBD between different concentrations of muscarine shows a significantly larger PBD at 3 and 10 µM compared to 1 µM muscarine, but no significant difference between 3 µM and 10 µM (MWU test). (ii) Summary graph of the incidence of spiking during PPs elicited following the application of different muscarine concentrations. Significant increase in the incidence of spikes during PPs in 3 and 10 µM muscarine compared to control conditions, but no incidents of spiking PPs with 1 µM muscarine. (WSR test, 1 µM: *n* = 10; 3 µM: *n* = 9; 10 µM: *n* = 10). Comparison of spiking during PPs shows a significant difference in spiking between both 3 and 10 µM compared to 1 µM, but no significant difference in spiking between 3 µM and 10 µM muscarine (MWU test). Continuous horizontal bars above the time-course plots indicate the extracellular presence in aCSF of the stated compounds in this and all following figures.

Muscarine elicited spiking PPs most consistently at 10 µM (**Figure 2D**; 10/10 cells) compared to 3 µM (6/9 cells) and 1 µM (0/11 cells). For this reason, we used 10 µM muscarine as the standard concentration to elicit PPs in all the following experiments (**Figures 3-7, Suppl. Figures 2-5**). During long-lasting muscarine applications, the induced depolarisation (PDB) and spiking declined over time, but declined less with 10 µM than with 3 µM (**Figure 2B**).

**Figure 3.**
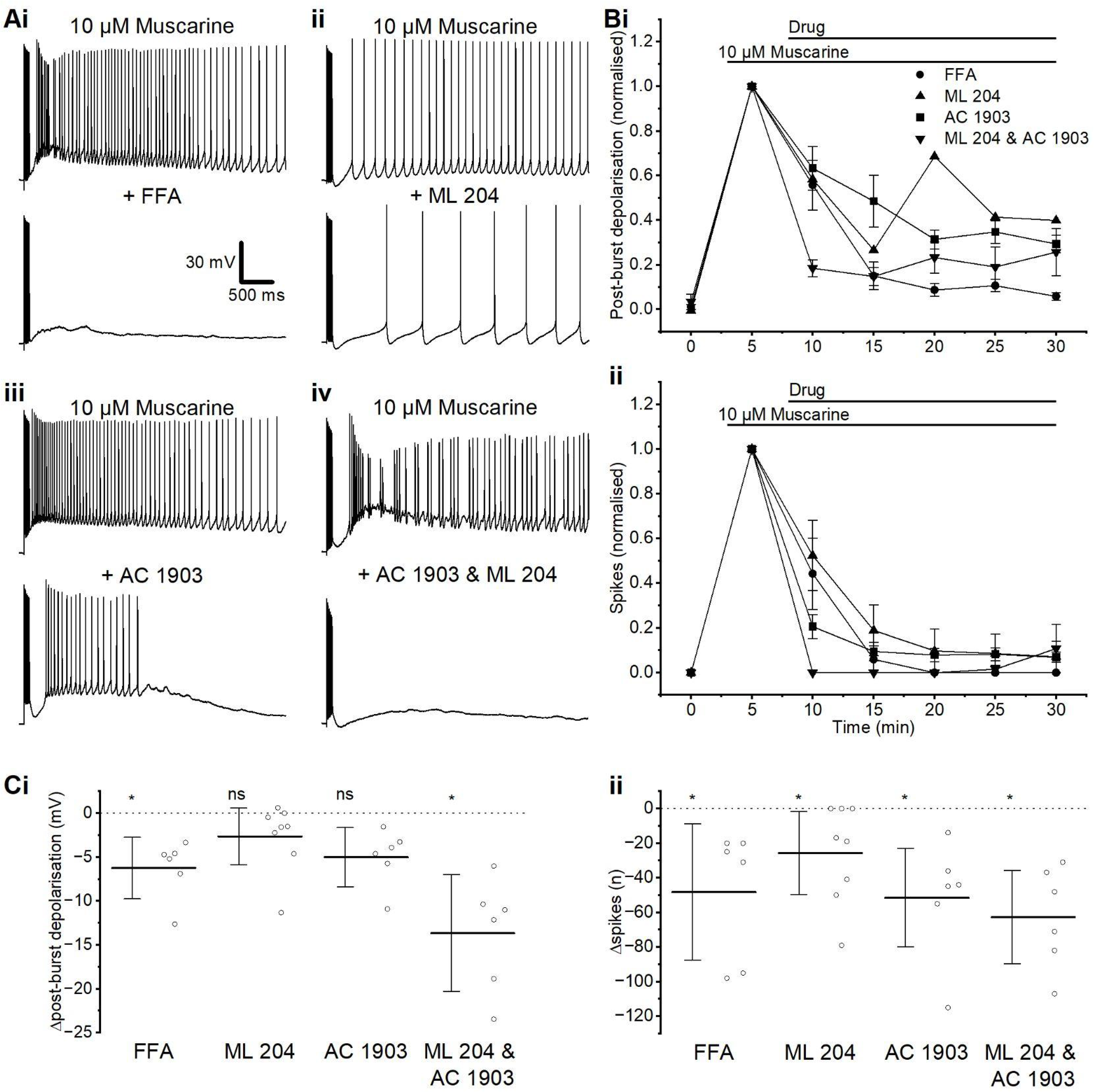
Evidence that plateau potentials depend on TRPC channels. **A** - Example traces showing the muscarinic PP before and after the application of TRP-blocking drugs FFA (i), ML 204 (ii), AC 1903 (iii), and ML 204 & AC 1903 (iv). **B** - Time course comparison of PBD (i) and spikes (ii) with different TRPC channel-blocking drugs. **C** - Summary plots of the PBD (i) and spikes (ii) before and after the application of different TRPC channel-blocking drugs. Significance was calculated in comparison to time-matched control measures. (i) A significant change in PBD was observed after the wash-in of FFA and ML 204 & AC 1903. (MWU test, FFA: *n* = 6; ML 204 & AC 1903: *n* = 6). There was no significant change in PBD when ML 204 or AC 1903 was washed in individually. (MWU test, ML 204; *n* = 8; AC 1903: *n* = 6). (ii) A significant change in spike count during PPs was observed in all groups compared with controls. (MWU test, FFA: *n* = 6; ML 204: *n* = 8; AC 1903: *n* = 6; ML 204 & AC 1903: *n* = 6).

Whilst these PPs were clearly mAChR-dependent, PPs have been observed elsewhere to be elicited following activation of several different metabotropic receptor types (Hagger-Vaughan & Storm, 2019; Perrier & Hounsgaard, 2003; Prager et al., 2020). To test whether metabotropic glutamate receptors (mGluRs) also can induce PPs in L2/3PCs, we applied tACPD, a selective agonist of the mGluR group I/II GluRs. After application of 15 µM tACPD, the spike-train protocol elicited a robust and reliable PP (**Suppl. Figure 1A**), with a significant PBD (control: -0.41 ± 0.10 mV; tACPD: 4.6 ± 1.1 mV, *n* = 12, *p* < 0.01) (**Suppl. Figure 1Bi**) and spikes (control: 0 ± 0 spikes; tACPD: 23 ± 7.4 spikes, *n* = 12, *p* < 0.01) (**Suppl. Figure 1Bii**). Since mAChRs and mGluRs are known to trigger similar signalling cascades, our results suggest that similar mechanisms of PP generation may be induced by a range or combination of different neuromodulators.

### L2/3PC PPs were dependent on TRPC channels

PPs have been shown to be dependent on transient receptor potential (TRP) channels in a variety of cell types (Hagger-Vaughan & Storm, 2019; Yan et al., 2009), and TRP channel activity is known to be enhanced by mAChR activation (Yoshida et al., 2012). Using the non-specific TRP channel blocker flufenamic acid (FFA; 50 µM) following the induction of PPs with muscarine, we observed a strong reduction in the PP relative to the time-dependant run-down in the control both in PBD (control: -1.8 ± 0.96 mV, *n* = 10; FFA: -6.2 ± 1.4 mV, *n* = 6, *p* = 0.02) and spikes (control: 5.4 ± 9.8 spikes, *n* = 10; FFA: -48 ± 15 spikes, *n* = 6, *p* = 0.01) (**Figure 3Ai & C**).

Previous work has identified the TRP channel subtypes TRPC4 and TRPC5 as being particularly abundant in the PFC (Fowler et al., 2007). We therefore used specific blockers for these channel subtypes to test if they were responsible for the muscarinic PP in L2/3PCs. Both the TRPC4 blocker ML 204 (10 µM) and the TRPC5 blocker AC 1903 (30 µM) produced a partial block of the PP (**Figure 3Aii & iii**), whilst the simultaneous application of both these blockers strongly reduced the PP (**Figure 3Aiv**). This suggests that both TRPC4 and TRPC5 are present in these cells and are involved in the generation of muscarine-induced PPs.

The change in mean PBD amplitude that we observed with the TRPC4 blocker ML 204 alone was not statistically significant compared to time-matched control experiments (control: -1.8 ± 0.96 mV, *n* = 10; ML 204: -2.6 ± 1.4 mV, *n* = 8, *p* = 0.83) (**Figure 3Ci**), but the spike number change relative to the control was significant (control: 5.4 ± 9.8 spikes, *n* = 10; ML 204: -26 ± 10 spikes, *n* = 8, *p* = 0.04) (**Figure 3Cii**). Also the TRPC5 blocker AC 1903 application gave a non-significant decrease in PBD (control: -1.8 ± 0.96 mV, *n* = 10, AC 1903: 5.0 ± 1.3 mV, *n* = 6, *p* = 0.12) (**Figure 3Ci**), but a significant decrease in spikes (control: 5.4 ± 9.8 spikes, *n* = 10; AC 1903: -52 ± 14 spikes, *n* = 6, *p* < 0.01) (**Figure 3Cii**).

The combination of ML 204 and AC 1903 gave the largest blocking effect on both change in PBD (control: -1.8 ± 0.96 mV, *n* = 10; ML 204 & AC 1903: -13.6 ± 2.6 mV (*n* = 6), *p* < 0.01) (**Figure 3Ci**) and post-burst spiking (control: 5.4 ± 9.8 spikes, *n* = 10; ML 204 & AC 1903: -63 ± 12 spikes, *n* = 6, *p* = 0.002 (**Figure 3Cii**).

Previous work has shown that TRPC channel modulation alters the impact of proximal apical EPSPs on PPs, but not the impact of distal apical EPSPs, and that FFA can be applied locally to probe the subcellular location of these channels (Lin et al., 2017). We attempted to determine the subcellular location of the PP-generating channels in L2/3PCs, by local pressure-application of FFA-containing aCSF from a glass pipette directed either towards the perisomatic region or distal apical dendrite, after PPs had been induced by bath-application of muscarine (**Suppl. Figure 2**). The PBD and the evoked spike number were clearly reduced in some cells following local application of FFA aimed at the perisomatic region, but not when applying FFA towards the dendrite (**Suppl. Figure 2A, i-ii**). There were also small changes in the mean values of these parameters across all cells tested only for FFA-applications towards the perisomatic region, but not towards the apical dendrite (**Suppl. Figure 2B i-ii**). However, the results varied considerably between cells, and neither the changes in PBD (control: -5.7 ± 1.5 mV, *n* = 6; FFA: -5.9 ± 0.58 mV, *n* = 10, *p* > 0.99) nor spikes (control: -30 ± 8.7 spikes, *n* = 6; FFA: -45 ± 12.4 spikes, *n* = 10, *p* = 0.45) were found to be statistically significant across all the tested cells. We are not sure why we found such a cell-to-cell variability and no statistically significant soma vs. dendrite differences across cells; several technical difficulties may have contributed to this.

### L2/3PC PPs were abolished by removal of extracellular calcium, but not by application of voltage-gated calcium channel (VGCC) blockers

To investigate whether the PPs are calcium-dependent (Oda et al., 2014; Owsianik et al., 2006), we tested the generation of PPs in aCSF with a low concentration of free Ca^2+^, containing 0.1 mM Ca^2+^ and 0.2 mM EGTA. In this condition, the addition of muscarine did not induce spiking PPs (**Figure 4A**): there was no significant change in post-burst spiking (control: 0 ± 0 spikes, muscarine; 0 ± 0 spikes, *n* = 6) (**Figure 4Cii**), but a small, significant PBD (control: -0.57 ± 0.20 mV, *n* = 6; muscarine: 0.44 ± 0.37 mV, *n* = 6, *p* = 0.03) (**Figure 4Ci**).

**Figure 4.**
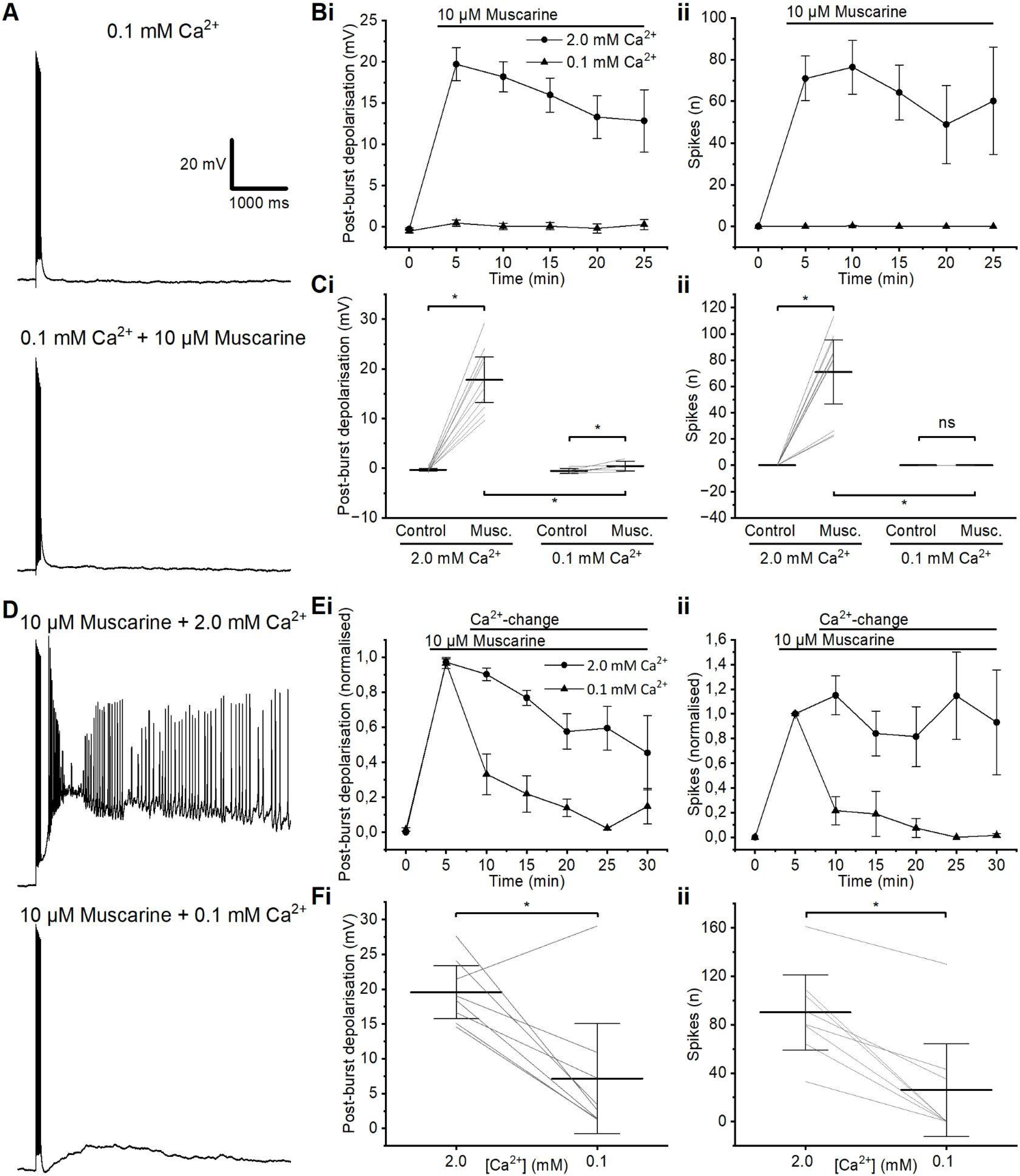
The plateau potential is Ca^2+^-dependent. **A** - Example traces of the effect of adding muscarine to low-Ca^2+^ aCSF, and the absence of an evoked PP. **B** - Time course summary of PBD (i) and spikes (ii) evoked by muscarine in normal- or low-Ca^2+^ aCSF. **C** - Summary graphs showing the effect of muscarine on PBD (i) and spikes (ii) in either normal- or low-Ca^2+^ aCSF. (i) Wash-in of 10 µM muscarine in both normal- and low-Ca^2+^ aCSF resulted in a significant change in PBD. (MWU test, 2.0 mM Ca^2+^: *n* = 10; 0.1 Ca^2+^: *n* = 6). (ii) A significant change in post-burst spiking was observed after the wash-in of muscarine in high Ca^2+^, but no spiking PPs in low Ca^2+^. (MWU test, 2.0 mM Ca^2+^: *n* = 10; 0.1 Ca^2+^: *n* = 6). There was a significant difference in both PBD and spike number between muscarine trials in normal- and low-Ca^2+^ aCSF (MWU test, *n* = 6). **D** - Example traces showing the effect of switching from normal-to low-Ca^2+^ aCSF on a muscarinic PP. **E** - Time course summary of PBD (i) and spikes (ii) evoked by muscarine and the subsequent change when switching from normal-to low-Ca^2+^ aCSF. Presented alongside control data with no change in Ca^2+^. **F** - Summary graphs showing the effect of switching from normal-to low-Ca^2+^ aCSF on PBD (i) and spikes (ii) in the presence of muscarine. A significant change in both PBD and spikes was observed after switching from a normal-Ca^2+^ aCSF to a low-Ca^2+^ aCSF (WSR, *n* = 8).

In normal aCSF containing 2.0 mM Ca^2+^, PPs were induced by muscarine, but switching to muscarine-containing, low-Ca^2+^ aCSF abolished the PPs (**Figure 4D**). The decrease in PBD (normal-Ca^2+^: 19.6 ± 1.6 mV; low-Ca^2+^: 7.2 ± 3.4 mV, *n* = 8, *p* = 0.02) and spikes (normal-Ca^2+^: 90 ± 13 spikes; low-Ca^2+^: 26 ± 16 spikes, *n* = 8, *p* = 0.01) after switching to low-Ca^2+^ aCSF was significantly different (PBD: *p* < 0.01; spikes: *p* < 0.01) from the changes seen in time-matched control cells when the aCSF was not changed during the experiment (**Figure 4Fi&ii**), indicating that these changes depended on extracellular Ca^2+^, and were not merely time-dependent.

In addition to the general Ca^2+^-dependence of PPs, we were interested in where in the cell the PP-generating Ca^2+^ influx occurred. Recording with low-Ca^2+^ aCSF (0.1 mM Ca^2+^, 0.2 mM EGTA, 4 mM Mg^2+^) in the bath, high-Ca^2+^ aCSF (5 mM Ca^2+^, 0.5 mM Mg^2+^) was pressure-applied (“puffed”) locally, via a glass pipette, to either the perisomatic or the distal, apical dendritic part of the cell in the presence of 10 μM muscarine (**Figure 5A**). In low-Ca^2+^ aCSF, with the presence of muscarine, PPs were rarely induced by the spike train protocol, but when Ca^2+^ was locally applied to the perisomatic region (while the slice bath was still perfused with low-Ca^2+^), a PP was evoked regularly (**Figure 5Bi & D**).

**Figure 5.**
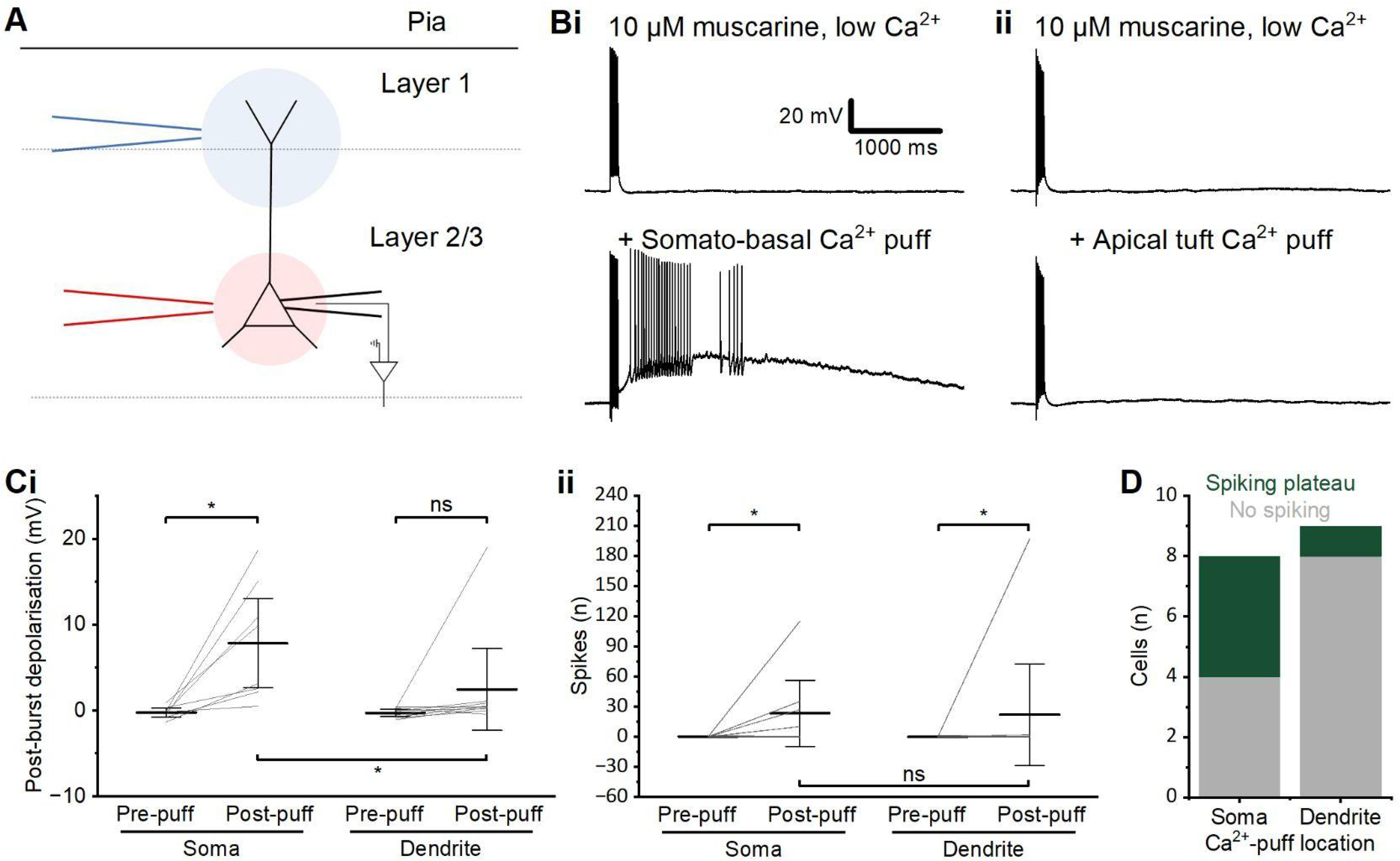
Peri-somatic/basal Ca^2+^ entry is essential for plateau potentials. **A** - Schematic representation of the experimental set-up for application of local Ca^2+^ either close to the soma (red) or apical dendrite (blue). **B** - Example traces showing the effect of local Ca^2+^ application either close to the soma (i) or apical dendrite (ii) in low-Ca^2+^ aCSF containing muscarine. **C** - Summary graphs showing the PBD (i) and spikes (ii) before and after calcium puffing at either the soma or dendrite. (i) A significant change in the PBD was observed after focal application of Ca^2+^ to the perisomatic region, but not after application to the apical dendrite (WSR, soma: *n* = 8; dendrite: *n* = 9). The post-puff PBD was significantly larger after the application of Ca^2+^ to the perisomatic region than after application to the dendrite (*p* = 0.018 by MWU). (ii) Post-burst spiking changed significantly in both application to the perisomatic region and to the dendrite (WSR, soma: *n* = 8; dendrite: *n* = 9), but not significantly different between perisomatic and dendritic application, largely due to one outlier cell that spiked following dendritic application (*p* = 0.22 by MWU). **D** - Summary graph of the incidence of spiking vs. non-spiking PPs evoked by different locations of Ca^2+^ application.

However, when Ca^2+^ was locally applied to the distal apical dendrite, only much weaker PBD, usually without spiking, were elicited, and far less reliably (**Figure 5Bii & D**). Thus, the PBD was significantly larger with perisomatic Ca^2+^ application than with dendritic Ca^2+^ application (perisomatic Ca^2+^: 7.8 ± 2.4 mV, *n* = 8; dendritic Ca^2+^: 2.5 ± 2.0 mV, *n* = 9, *p* = 0.01). Also, the prevalence of spiking PPs was much higher after a perisomatic Ca^2+^ puff (**Figure 5D**), i.e. PPs were seen far more often after perisomatic than after apical dendritic Ca^2+^ application. However, there was no statistically significant difference in post-burst spiking between the perisomatic-puff and dendrite-puff groups, because one cell in the latter group showed a large, spiking PP, whilst none of the others in the group did (perisomatic Ca^2+^-puff: 27 ± 16 spikes, *n* = 8; dendritic Ca^2+^-puff: 22 ± 22 spikes, *n* = 9, *p* = 0.22) (**Figure 5Ci**). Again, there might have been technical difficulties with the local application in some cases.

Since we found that the PPs were reduced by blockers of TRPC4 and TRPC5 channels (**Figure 3**), which are known to be Ca^2+^-permeable (Owsianik et al., 2006), and we also found that the PPS were Ca^2+^-dependent (**Figures 4 & 5**), it seems likely that Ca^2+^ entry through TRPC4 and TRPC5 channels was involved in PP generation. However, since PPs have been demonstrated to be dependent on voltage-gated Ca^2+^ channels (VGCCs) in other cell types (Fraser & MacVicar, 1996; Heng et al., 2011; Lo & Erzurumlu, 2002; Williams & Fletcher, 2019), we sought to determine whether VGCCs could also play a role here.

We found that the non-specific VGCC blocker cadmium (Cd, 100 μM) suppressed PPs (**Suppl. Figure 3Ai**), with large, significant reductions in both PBD (control: -1.8 ± 0.96 mV, *n* = 10; CdCl_2_: -10.9 ± 1.7 mV, *n* = 6, *p* < 0.01) (**Suppl. Figure 3Bi**) and spikes (control: 5.4 ± 9.8 spikes, *n* = 10; CdCl_2_: -95 ± 12 spikes, *n* = 6, *p* < 0.01) relative to the control (**Suppl. Figure 3Bii**).

Nickel (Ni^2+^), at 50 μM concentration, is known to block both T-type and R-type VGCCs (Magee & Johnston, 1995). We observed a range of effects of 50 μM Ni^2+^ on PPs (**Suppl. Figure 3Aii**), but found no significant differences in either PBD (control: -1.8 ± 0.96 mV, *n* = 10; NiCl_2_: -1,4 ± 1.6 mV, *n* = 5, *p* = 0.77) (**Suppl. Figure 3Bi**) or spiking (control: 5.4 ± 9.8 spikes, *n* = 10; NiCl_2_: -29 ± 16 spikes, *n* = 5, *p* = 0.11) compared to the control (**Suppl. Figure 3Bii**). Wash-in of the L-type VGCC blocker nifedipine (10 μM; **Suppl. Figure 3 Aiii**) also caused no significant change in either PBD (control: -1.8 ± 0.96 mV, *n* = 10; nifedipine: -2.0 ± 0.88 mV, *n* = 5, *p* = 0.77) (**Suppl. Figure 3Bi**) or spiking (control: 5.4 ± 9.8 spikes, *n* = 10; nifedipine: -18 ± 8.4 spikes, *n* = 5, *p* = 0.35) (**Suppl. Figure 3Bii**). Similarly, the P-type VGCC blocker PD-173212 (10 μM; **Suppl. Figure 3Aiv**) caused no significant changes in either PBD (control: -1.8 ± 0.96 mV, *n* = 10; PD-173212: -0.77 ± 1.1 mV, *n* = 7, *p* = 0.42)(**Suppl. Figure 3Bi**) or spiking (control: 5.4 ± 9.8 spikes, *n* = 10; PD-173212: -10 ± 8.9 spikes, *n* = 7, *p* = 0.35) (**Suppl. Figure 3Bii**).

Having established the importance of extracellular Ca^2+^ in generating PPs, we wished to investigate any possible involvement of intracellular Ca^2+^ stores. Intracellular Ca^2+^ release has been implicated in PPs (Mejia-Gervacio et al., 2004; Power & Sah, 2002) and involved in the insertion of TRPC5 channels into the cell membrane and other TRPC channel activation (Tai et al., 2011), leading to PPs. mAChRs can induce Ca^2+^ release from intracellular stores via the αGq pathway, causing activation of IP3 receptors (IP3Rs) in the endoplasmic reticulum.

To test whether IP3Rs are involved in PP-generation in L2/3 pyramidal cells, we included the IP3Rs blocker heparin in the intracellular recording solution (5 mg/ml), and allowed 20 minutes after achieving whole-cell configuration, for it to diffuse into the cell (**Suppl. Figure 4)**. Heparin had no apparent effect prior to muscarine application. Thus, we found no difference in the PBD (control ICS: -0.33 ± 0.12 mV, *n* = 10; heparin ICS: -0.21 ± 0.25 mV, *n* = 4, *p* > 0.999) or spiking (control ICS: 0 ± 0 spikes, *n* = 10; heparin ICS: 0 ± 0 spikes, *n* = 4) between the control and heparin groups, suggesting that heparin did not affect basal properties. However, after 20 min. of heparin “dialysis”, application of muscarine did not induce robust PPs (**Suppl. Figure 4A**), in sharp contrast to cells without i.c. heparin (**Figures 1-3; Suppl. Figure 4B-C**). Thus, although most heparin-filled cells showed an AHP reduction and a small PBD (mean PBD increased) after muscarine was applied (**Suppl. Figure 4A** and **Ci**), the changes were not statistically significant compared to time-matched recordings with control intracellular solution, neither for PBD (control: -0.21 ± 0.25 mV; muscarine: 1.1 ± 0.48 mV, *n* = 4, *p* = 0.25) (**Suppl. Figure 4Ci**) nor spikes (control: 0 ± 0 spikes; muscarine: 0 ± 0.0 spikes, *n* = 4) (**Suppl. Figure 4Cii**). However, the finding that intracellular heparin almost fully blocked the PP, suggests that Ca^2+^ release from intracellular stores is important for the generation of PPs in L2/3PCs.

### Plateau potentials could be generated in the absence of apical dendrites

Although plateau potentials have been found to depend on the apical dendrite in several cell types, and an apical dendritic origin is often assumed to be typical, there are reasons to think that PPs may also be generated in perisomatic or basal dendritic compartments in some cases. Thus, there is evidence that the TRPC channels that appear to be necessary for PPs in the L2/3 PCs (**Figure 3**) are expressed only in the perisomatic compartment (Fowler et al., 2007).

To test whether the PPs in the L2/3 PCs are generated in or depend on the apical dendrite, we disconnected the apical dendrites from the perisomatic compartments by a scalpel blade cut through the slice between the L2/3 PC somata and the pia (**Figure 6Ai**), to disconnect L1 and the superficial portion of L2 from the deep L2 and L3. Shortly (10-15 minutes) after the slice was cut, somatic whole-cell recordings were obtained, with Alexa- and biocytin-filled patch pipettes, from L2/3 PC somata. During and after the recording, the Alexa-filled cells were observed immediately with fluorescence microscopy, and the biocytin-stained cells were later reconstructed, thus confirming that the distal part of the apical dendrite was indeed disconnected from the perisomatic compartments, at distances ranging from 20 to 260 μm (**Figure 6Aii & iii Figure 7**).

**Figure 6.**
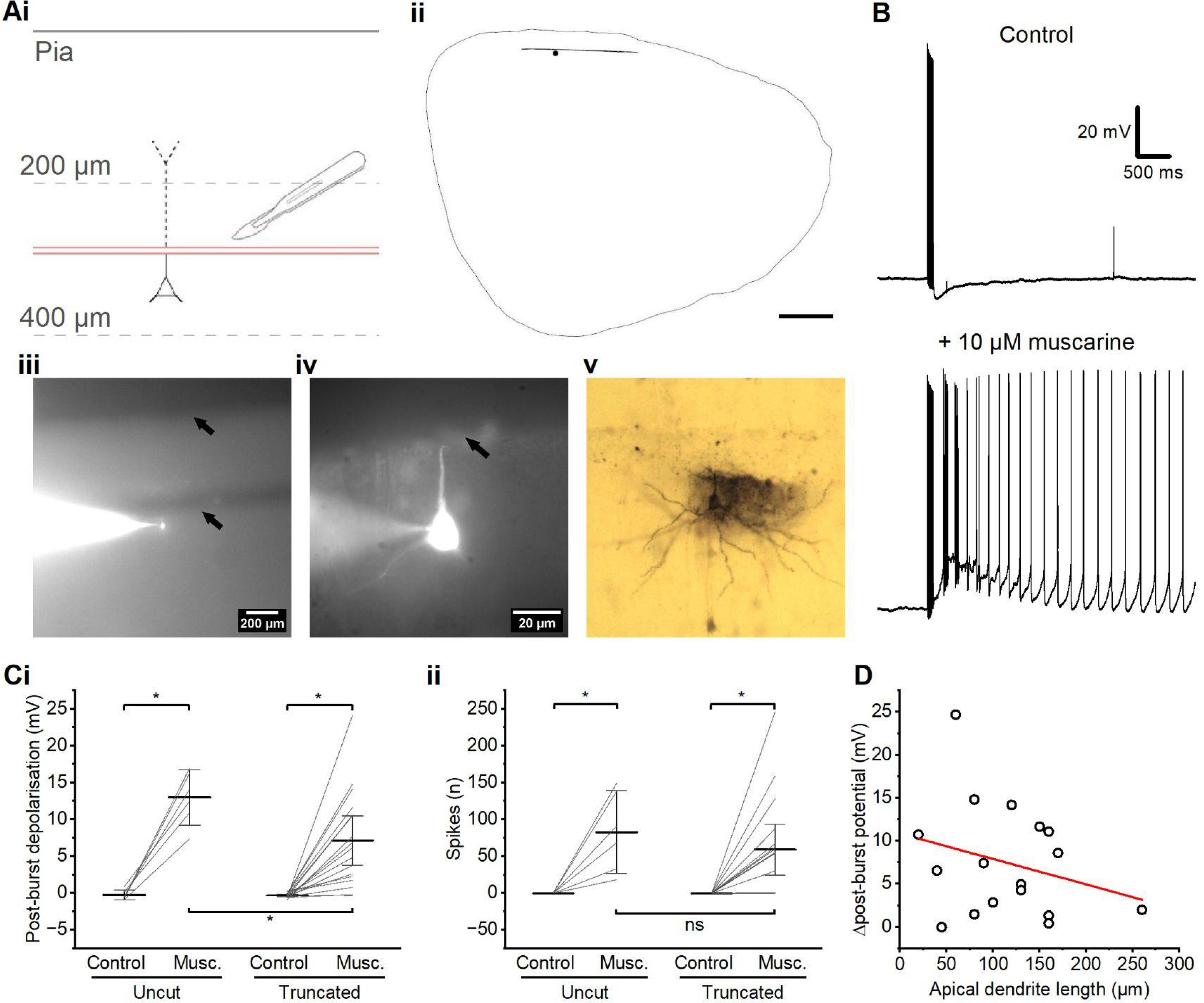
Plateau potentials can be elicited in the absence of the apical trunk. **A** - (i) Schematic illustration of experimental procedure for slice cutting, with the two red lines illustrating the cut, and the dotted lines illustrating the disconnected distal apical dendrite. (ii) Tracing of a coronal prefrontal rat brain slice, with a cut (illustrated by a black line), and a L2/3PC (represented by a black dot). (iii) A low magnification photomicrograph of the same cell illustrated in (ii), filled with Alexa 555. The top black arrow indicates the pia, while bottom the black arrow indicates the cut. (iv) A high magnification photomicrograph of the same cell as seen in (iii and iv), with a black arrow indicating the cut. (v) A photomicrograph of the same cell as in (iv) following filling with and staining for biocytin. **B** - Example traces of a muscarinic PP evoked in a cell with a truncated apical dendrite. **C** - Summary plots comparing the PBD (i) and spikes (ii) in cells with truncated and intact apical dendrites. (i) A significant change in PBD was observed after the wash-in of muscarine in both the control and truncated groups (WSR test, truncated dendrite; *n* = 17, control; *n* = 6). There was also a significant difference in PBD between the truncated and control groups after the wash-in of muscarine (*p* = 0.024 by MWU test). (ii) A significant change in spikes was also observed after the wash-in of muscarine in both groups (WSR test, truncated dendrite; *n* = 17, control; *n* = 6). No significant difference in spikes was observed between the two groups after the wash-in of muscarine (*p* = 0.19 by MWU test). **D** - Summary plot of change in PBD following muscarine application against measured remaining apical dendrite length post-cut. (Spearman’s ρ = -0.18; *n* = 17; *p* = 0.48).

**Figure 7.**
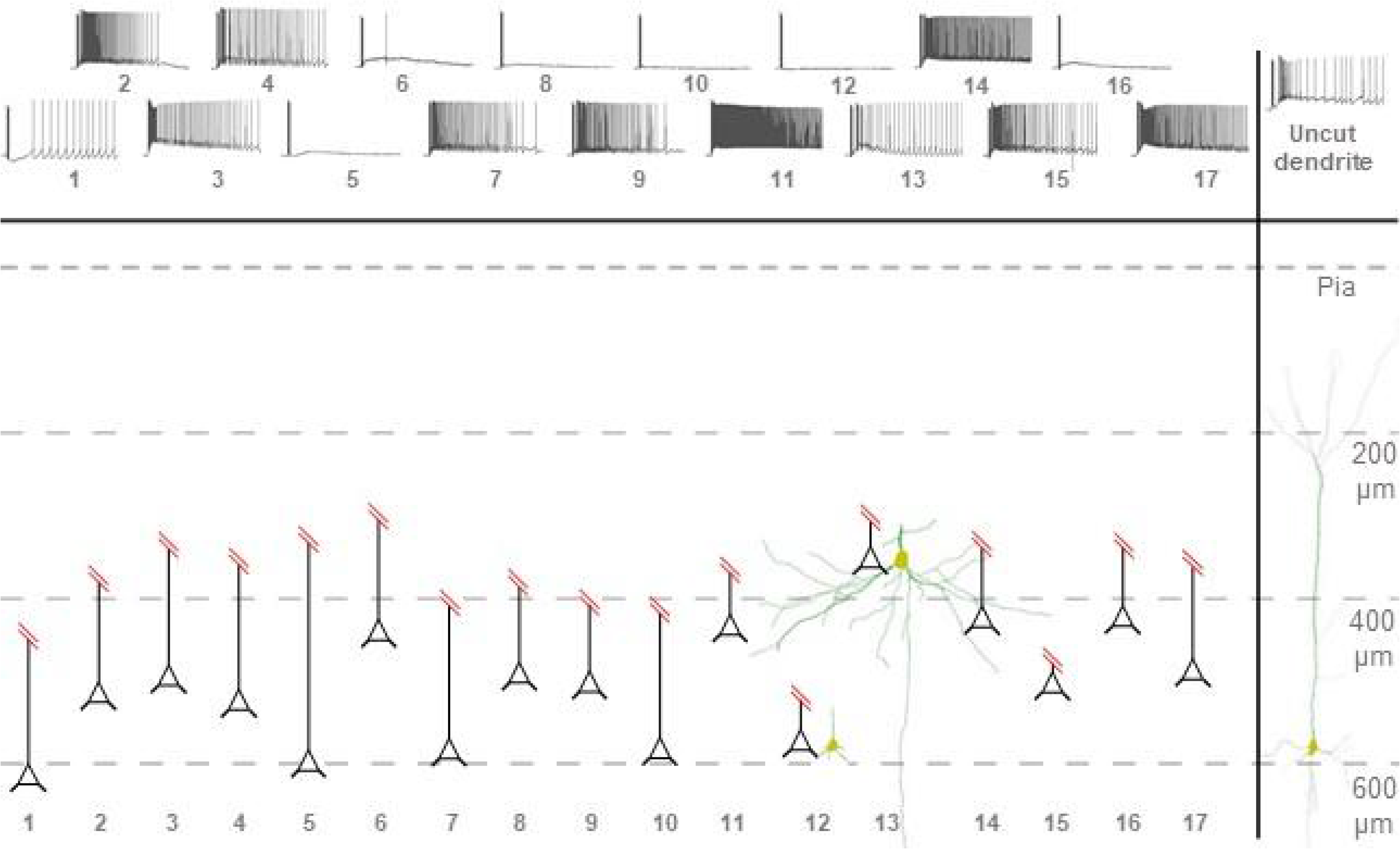
Overview of cells with truncated apical dendrites. Overview of cell morphologies post-cut as described in Figure 6. Schematic representations of cells represent soma distance from the pia mater (Pia) and intact apical dendrite trunk. Two reconstructed morphologies of truncated cells and one of a control cell are also included for further context. Example traces after the wash-in of 10 µM muscarine are included in the top row with cell numbering.

No clear differences in input resistance or resting membrane potential between truncated (cut) cella and control cells were found two minutes after break-in (data not shown). However, truncated cells that did not show spiking PPs after the wash-in of 10 µM muscarine had a tendency to require a smaller, positive holding current to hold the membrane potential at -60 mV before the spike-burst protocol at time = 0 min (data not shown).

In the truncated cells, 10 µM muscarine induced PPs that appeared quite similar to those in intact cells (**Figure 6B**). Thus, most of the truncated cells showed PBDs that were significantly increased by muscarine (control: -0.29 ± 0.08 mV; muscarine: 7.2 ± 1.6 mV, *n* = 17, *p* < 0.01), but the muscarinic PPs/PBD amplitudes were smaller than in control cells (cut with muscarine: 7.2 ± 1.6 mV, *n* = 17; uncut with muscarine: 13.0 ± 1.5 mV, *n* = 6, *p* = 0.02) (**Figure 6Ci**). Muscarine also induced PP-evoked post-burst spiking in truncated cells (control: 0.1 ± 0.1 spikes; muscarine: 59 ± 16 spikes, *n* = 17, *p* < 0.01), with no significant difference between cut (truncated) and uncut cells (cut + muscarine: 59 ± 16 spikes, *n* = 17; uncut + muscarine: 83 ± 22 spikes, n = 6, *p* = 0.19) (**Figure 6Cii**).

Similar to uncut cells, truncated cells exhibited a range of PP types (**Suppl. Figure 5**), but the PBD amplitude was not correlated with the remaining apical dendrite length (Spearman’s ρ = -0.18, *n* = 17; *p* = 0.48) (**Figure 6D**).

These results (**Figure 6**), combined with our results from local applications of Ca^2+^ (**Figure 5**), indicate that muscarinic PPs in L2/3 PCVs can be generated in the absence of the distal apical dendrite, thus probably being generated by persomatic- and/or basal dendritic-dependent mechanisms.

### Optogenetic activation of cholinergic afferents

Having shown that bath-application of muscarine paired with a somatic spike-train can regularly induce spiking PPs in L2/3PCs, we wanted to investigate whether we could elicit spiking PPs in this cell type also by pairing somatic spike-trains with release of endogenous ACh from local cholinergic neurons and surviving cholinergic axons with presynaptic terminals in the slice (Obermayer et al. (2018). To do this, we used PFC brain slices from mice expressing both channelrhodopsin-2 (ChR2) and enhanced yellow fluorescent protein (EYFP) in cells expressing choline transferase (ChAT), i.e. cholinergic interneuron (Granger et al., 2020; Hedrick & Waters, 2015; Obermayer et al., 2019) (**Suppl. Figure 5A**). After obtaining somatic whole-cell recording from an EYFP-labelled (i.e. cholinergic) neuron (n=5), the slice was flashed with trains of brief (20 ms) light flashes at 25 Hz. Each flash reliably triggered an action potential in 4 of the 5 EYFP-labelled neurons tested (**Suppl. Figure 5B**), thus demonstrating that the neurons were activated by the flashes.

Next, we tested the effect of light-induced release of endogenous ACh on whole-cell recorded L2/3PCs (*n* = 17). In these cells, a train of five or ten light flashes at 25 Hz, (similar to the protocol used by Obermayer et al (2018) (**Suppl. Figure 5C**), induced a small but significant PBD after a current-evoked burst of 7 spikes, but the PBD did not trigger spiking. Thus, there was a significant difference in post-burst membrane potential between the unpaired spike-trains and the spike-trains paired with light stimulation (unpaired: -0.15 ± 0.11 mV; with light stimulation: 0.69 ± 0.32 mV, *n* = 17, *p* = 0.01) (**Suppl. Figure 5D**).

## Discussion

### Variable properties of muscarine-dependent plateau potentials

In this study, we examined the effect of the metabotropic cholinergic agonist muscarine on the membrane potential of PFC L2/3PCs following a train of action potentials evoked by somatic injection of brief current pulses. In control conditions, all cells tested exhibited a medium-duration (∼0.2 s) afterhyperpolarization (AHP). Application of 10 µM muscarine abolished the AHP and usually led to a long-lasting (∼seconds) plateau potential (PP) that often triggered repetitive action potential firing. However, the PPs were variable. With 10 µM muscarine, 70% of PPs were accompanied by persistent firing (PF), i.e. spiking at a fairly constant rate that persisted for several seconds. Other PPs were self-terminating, with spiking that gradually slowed and stopped before the membrane potential returned to baseline; or there was only a large ADP with only a few, low-frequency action potentials. Finally, in around 30% of cells using 3 µM muscarine, only an ADP with no spiking was observed. This variability might be due to heterogeneity in the tested sample of cells, perhaps reflecting subsets of cells with different sets of channels or receptors (Karimi et al., 2020; Luo et al., 2017; Weiler et al., 2023). The probability of generating a spiking PP was also greatly dependent on the concentration of muscarine used (**Figure 2C**).

### Evidence that plateau potentials depend on TRPC channels

Based on our results with various TRPC channel agonists and antagonists, we concluded that both TRPC4 and 5 were important for the generation of PPs.

TRPC5 currents can be activated or enhanced in different ways: intracellular signalling from G-protein coupled receptors can enhance TRPC currents via direct interaction with Gi proteins (Jeon et al., 2012); translocation of vesicular TRPC channels to the membrane leads to TRPC5 currents and PPs (Hong et al., 2012; Tai et al., 2010); and release of Ca^2+^ from intracellular stores increases TRPC current amplitudes through the channels interaction via STIM1 (Pani et al., 2012).

Whether or not TRPC channels underlie muscarinic ADPs and PPs may be highly cell-specific, even within specific brain regions. Thus, previous work has both found and failed to find evidence that TRPC5 channels contribute to muscarinic ADPs in various L5PCs in the PFC of rodents (Dasari et al., 2013; Yan et al., 2009). Whilst our results seemed to show a greater effect on PPs using a TRPC5 antagonist than a TRPC4 antagonist, we did not test a range of concentrations, so the concentrations of TRPC4 and TRPC5 antagonists used might not have been strictly comparable. Therefore we cannot conclude which of these channel types is most important for the PPs in these cells, although both seem to contribute. Further work is needed to determine the exact contributions of different TRPC subtypes to these PPs in these cells.

### Evidence that plateau potentials depend on Ca^2+^ but not on voltage-gated Ca^2+^ channels

TRPC channels are activated by intracellular Ca^2+^ ions (Hasan & Zhang, 2018), which may enter the cell via channels in the plasma membrane or may be released from intracellular Ca^2+^ stores. Our observations that PPs were inhibited by Cd^2+^, but not by specific VGCC blockers, suggest that the PPs were generated by TRPC channels activated by intracellular Ca^2+^ that entered the cell via another route than VGCC.

Calcium release-activated channels (CRAC) channels such as ORAI channels have been shown to be blocked by Cd^2+^ with an IC50 of 200 μM (Hoth & Penner, 1993), and ORAI channels and TRPC channels have been shown to associate in complexes (Liao et al., 2009). Thus, the plateau potentials (PPs) may be generated by TRPC channels that form complexes with ORAI-based CRAC channels, whose Ca^2+^ influx activates the TRPC channels.

This mechanism of PP generation would be markedly different from that of the “typical” dendritic calcium-plateau/-spike in the distal apical hot zone in L5 PCs, where strong depolarisation is required to activate L-type (Heng et al., 2011; Simon et al., 2003) or R-type (Kuzmiski & MacVicar, 2001; Williams & Fletcher, 2019) VGCCs. In L2/3PCs in the human neocortex, dendritic action potentials caused by coincident input have been shown to be Ca^2+^-based and often induce somatic spiking (Gidon et al., 2020).

### Muscarinic plateau potentials in L2/3PCs are *non-apical*: they were not eliminated by truncation of the apical dendrites

PPs in cortical pyramidal cells have previously been suggested to be largely of dendritic origin, often triggered by convergent excitatory input to somatic and apical compartments (H. Takahashi & Magee, 2009). There is also evidence that apical dendritic PPs in response to convergent excitatory input is important for both reporetable sensory perception (Suzuki & Larkum, 2020; N. Takahashi et al., 2016)) and in the overall alert brain state during normal wakefulness *in vivo* (Suzuki & Larkum, 2020; N. Takahashi et al., 2016), thus probably being involved in at least some forms of access consciousness (Suzuki & Larkum, 2020; N. Takahashi et al., 2016).

To test whether the muscarinic plateau potentials in L2/3 PCs also depend on apical dendritic activity, we physically cut the apical dendrite at various distances from the soma. We found that the somatically recorded plateau potentials still persisted and triggered spiking (PF) even when most of the apical dendrite was disconnected (**Figure 6**). In addition, when the PPs were eliminated by perfusing the slice chamber with a low-Ca^2+^ aCSF, local application of Ca^2+^ only to the perisomatic region restored PP generation (**Figure 5**). Hence, it seems that these muscarinic plateau potentials are generated largely in the perisomatic and basal parts of the L2/3 PCs including basal dendrites and soma (and possibly even parts of the axon, although the latter seems less likely). PPs evoked by basal dendritic input, and associated Ca^2+^ transients occurring in the basal dendrites, have been found previously in L5PCs in the prefrontal cortex (Milojkovic et al., 2007). However, it remains to be seen whether such PPs can be generated in vivo by natural basal synaptic input alone.

The well-known dendritic Ca^2+^ spikes/plateaus of L5 PCs can typically be triggered by back-propagating Na^+^ action potentials originating in the soma, combined with local dendritic excitatory synaptic input. This activates VGCCs in the nexus of the apical trunk, the so-called Ca^2+^ hot zone, and the subsequent Ca^2+^ entry can trigger somatic/axonal action potential bursts (M. Larkum, 2013) This general mechanism may also occur in L2/3 cortical pyramidal cells. This study, however, demonstrates that the apical dendrite is not needed (at least not in its entirety) to initiate and maintain PPs and PF in some L2/3PCs.

### Consequences of amplification by plateau potentials in L2/3PCs

As described above, we found that cholinergic modulation caused L2/3PCs to greatly increase their spike output in response to a train of stimuli. The consequences of such amplification will be felt both locally and across large areas of the neocortex.

Thus, through the long-range cortico-cortical projections of L2/3PCs the increased output during cholinergic influence will impact remote cortical targets. Within the PFC, L2 and L3PCs send axons to L5PCs (Cheriyan & Sheets, 2018), and specifically to cortico-thalamic PCs (Collins et al., 2018) which target thalamic nuclei, thereby completing a multi-synaptic reciprocal loop. Outside the PFC, L2 and L3PCs axons project primarily to other cortical areas, including the contralateral PFC, but also to subcortical areas such as the baso-lateral amygdala (BLA), forming direct reciprocal connections, and the striatum (Anastasiades et al., 2019).

The origin of inputs to these cells may indicate which signals will be amplified by PPs in L2/3PCs. Layer 3 PCs receive their main excitatory input from the mediodorsal thalamus (MDT) (Collins et al., 2018), which is known to be important for attention and memory (Golden et al., 2016), with PFC-MDT connections being essential for object (Pergola et al., 2013) and recency (Cross et al., 2012) recognition. Layer 2 PCs also receive input from the BLA (Little & Carter, 2013). BLA activity and reciprocal connections between the BLA and PFC have been shown to be important in encoding emotional states, and in particular emotionally weighted memories (Burgos-Robles et al., 2017).

### Functional consequences of the non-apical origin of the plateau potentials

The non-apical origin of these PPs suggest that they preferentially will boost excitatory input of perisomatic and basal dendritic origin, as opposed to apical dendritic input, unlike the apical dendritic PPs and Ca^2+^-spikes typically observed in L5PCs (M. E. Larkum et al., 1999). Thus, the non-apical PPs of rat L2/3PCs seem poorly suited to perform the key type of coincidence detection and dendritic integration between two different streams of information that has been thoroughly characterised in rodent L5PCs (M. Larkum, 2013; M. E. Larkum et al., 1999; Major et al., 2013; Phillips et al., 2016), and which forms the core of the dendritic integration theory of consciousness (DIT; (Aru et al., 2020). An interesting, still open question is whether L2/3PCs in humans and other primates, whose apical dendrites are far longer and equipped with different ionic conductances than in rodents (Kalmbach et al., 2018), and function more like rodent L5PCs (Beaulieu-Laroche et al., 2018) than rodent L2/3PCs in this respect. In other words: do human L2/3 PCs have an apical dendritic PP mechanism that allows nonlinear integration between different information streams, like in rodent L5PCs?

### Effects of bath-applied muscarine vs. optogenetically released ACh

In most of our experiments, we used bath application of muscarine in an attempt to probe the muscarinic effects of ACh release. However, the effects of bath application may of course differ from those of physiological ACh release. To better approximate natural ACh modulation, we activated cholinergic axons and terminals in the slice by optogenetics. This often caused a small, transient PBD, but no clear, persistent PPs or spiking. We do not know why the optogenetic method produced smaller PBD than bath-application, or whether the former effect was weaker (e.g. due to inferior function of the cholinergic axons and terminals in the slice) or the latter stronger than natural cholinergic modulation. However, our results do not seem to exclude the possibility that natural cholinergic modulation can induce PP generation in vivo, e.g. when combined with other modulators. Thus, although our optogenetic flash protocol evoked action potentials in the soma of local cholinergic interneurons (**Suppl. Figure 5B**), we do not know if it efficiently triggered action potentials in long-range cholinergic axons and presynaptic terminals in the slice, axons that were cut off from basal the forebrain cholinergic nuclei that provide the majority of the ACh input to the PFC in vivo (Ballinger et al., 2016). Perhaps many of the cut cholinergic axons were not excited or unable to release ACh, thus reducing the flash-induced release to a fraction of that occurring in vivo. In addition, cortical neurons in vivo are normally exposed to a cocktail of neuromodulators, which can sometimes have strongly synergistic effects. Several studies have shown that low concentrations of two neuromodulators applied together can induce PPs, even when one modulator alone has far less effect (Hagger-Vaughan & Storm, 2019; Park & Spruston, 2012). In the PFC L2/3PCs, we found that also mGluR activation induced PPs. Therefore it is plausible that these cells can exhibit PPs in vivo when receiving natural, diverse neuromodulatory inputs. Experiments using combinations of agonists and optogenetically induced neuromodulator release, preferentially in vivo, are needed to further test these possibilities.

### Summary

We found that layer 2/3 pyramidal neurons in rat prefrontal cortex, when modulated by muscarinic cholinergic (mAChRs) input can generate plateau potentials (PPs) with sustained spiking, and that this can occur independently of distal apical dendrites.

The plateaus seem to depend on calcium, both external Ca^2+^ and internal Ca^2+^ stores, and on TRPC4 and TRPC5 cation channels, but not on voltage-gated Ca^2+^ channels. Since L2/3PCs have long-range cortical and subcortical connections, the increased, persistent spiking activity caused by muscarinic modulation of these cells, may have a widespread impact in brain states with high cholinergic input to the neocortex, such as during REM sleep and wakefulness.

## Figures

**Supplementary figure 1.**
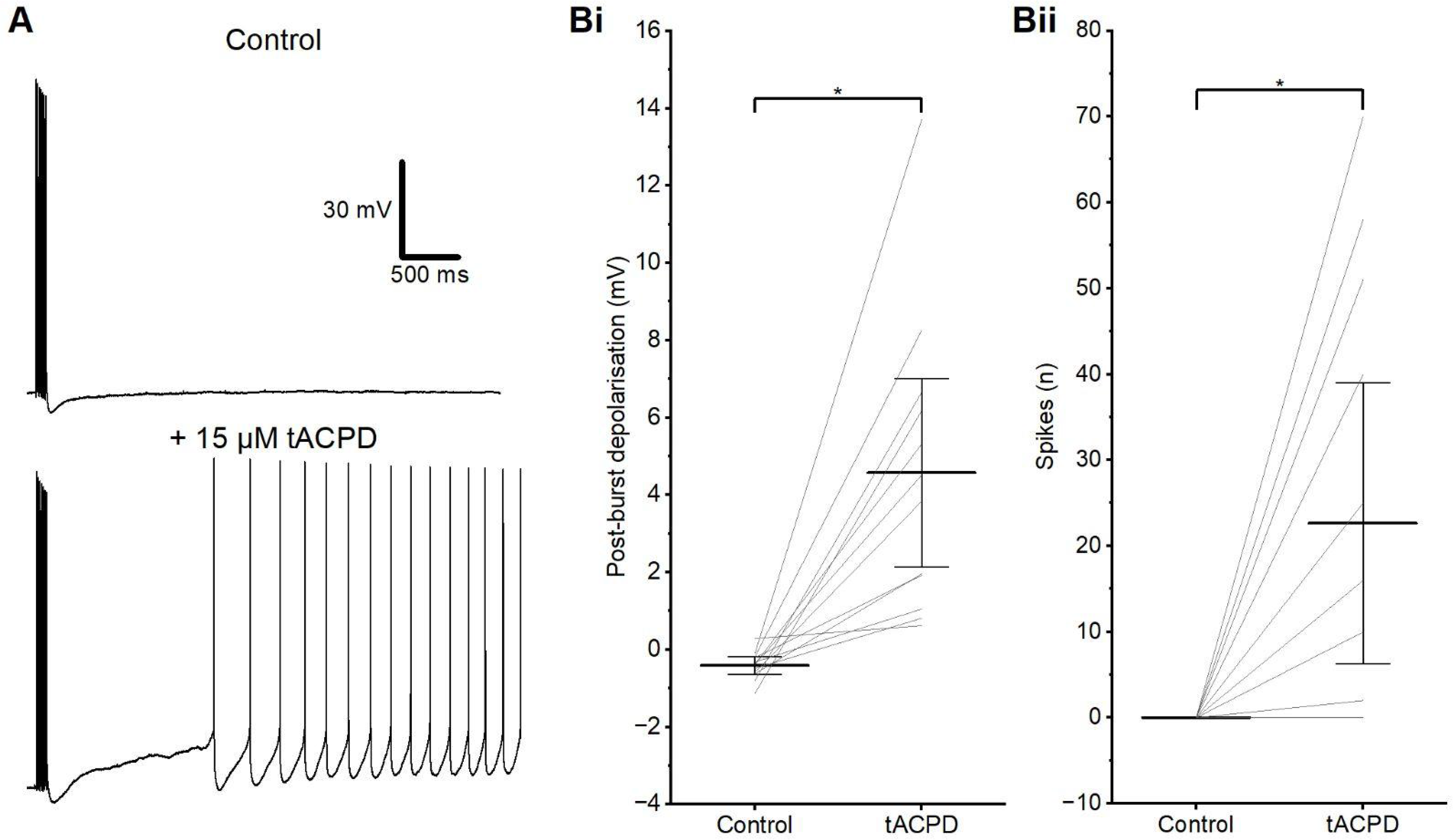
Plateau potentials can be induced by mGlu receptors. **A** - Example traces showing the PP induced by tACPD. **B** - Summary plots of the PBD (i) and spikes (ii) following the application of tACPD. Both PBD (i) and post-burst spiking (ii) change significantly after wash-in of tACPD (WSR test, *n* = 12).

**Supplementary figure 2.**
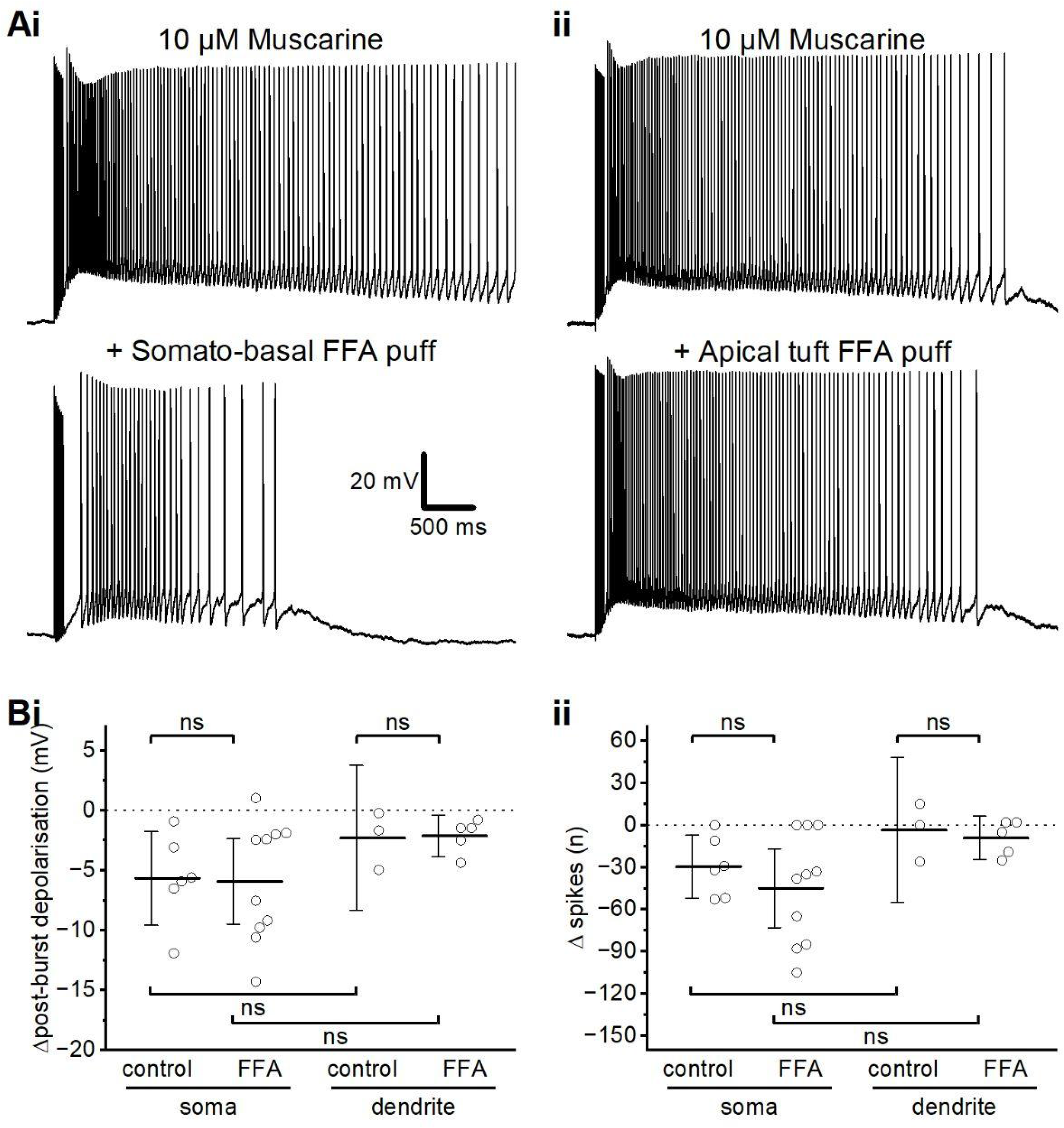
Local FFA application at the soma and apical dendrite. **A** - Example traces showing the PP before and after local application of FFA (200 μM in the puffing pipette) aimed at the perisomatic region (i) or the apical dendrite (ii). **B** - Summary plots of the evoked PBD (i) and change in the number of evoked spikes (ii) before and after pressure-application of normal aCSF (Control) or FFA-containing aCSF aimed at the perisomatic region or apical dendrite. Although both the PBD and the evoked spike number were clearly reduced in some cells following application of FFA towards the perisomatic region, but not when applying FFA towards the dendrite, as illustrated in **Ai-ii**, and there were small changes in the mean values of these parameters across all cells tested only for FFA-applications towards the perisomatic region, not towards the dendrite (**Ai-ii**), the results varied considerably between cells, and neither the changes in PBD nor spikes weres found to be statistically significant across all the tested cells (MWU test, somatic control: *n* = 6; somatic FFA: *n* = 10; dendritic control: *n* = 3; somatic FFA: *n* = 5).

**Supplementary figure 3.**
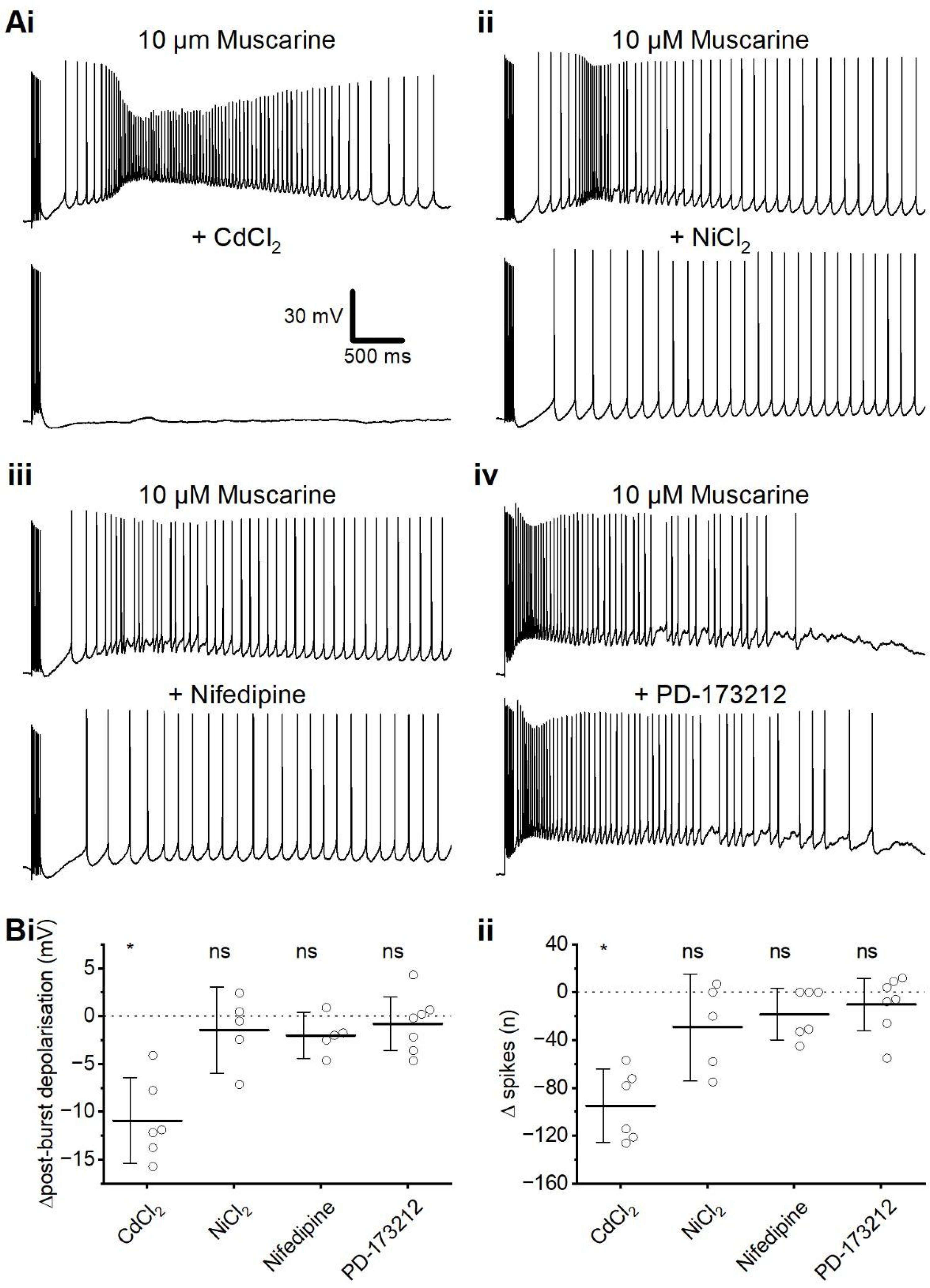
PPs are not prevented by VGCC blockers. **A** - Example traces of muscarinic PPs before and after application of VGCC blockers CdCl_2_(i), NiCl_2_ (ii), nifedipine (iii), and PD-173212 (iv). **B** - Summary plots of the change in PBD (i) and spikes (ii) after the application of different VGCC-blocking drugs. A significant change in PBD and post-burst spiking was observed after the wash-in of CdCl_2_, but not after the wash-in of the other VGCC blockers (MWU test, CdCl_2_: *n* = 6; NiCl_2_: *n* = 5; Nifedipine: *n* = 5; PD-173212: *n* = 7).

**Supplementary figure 4.**
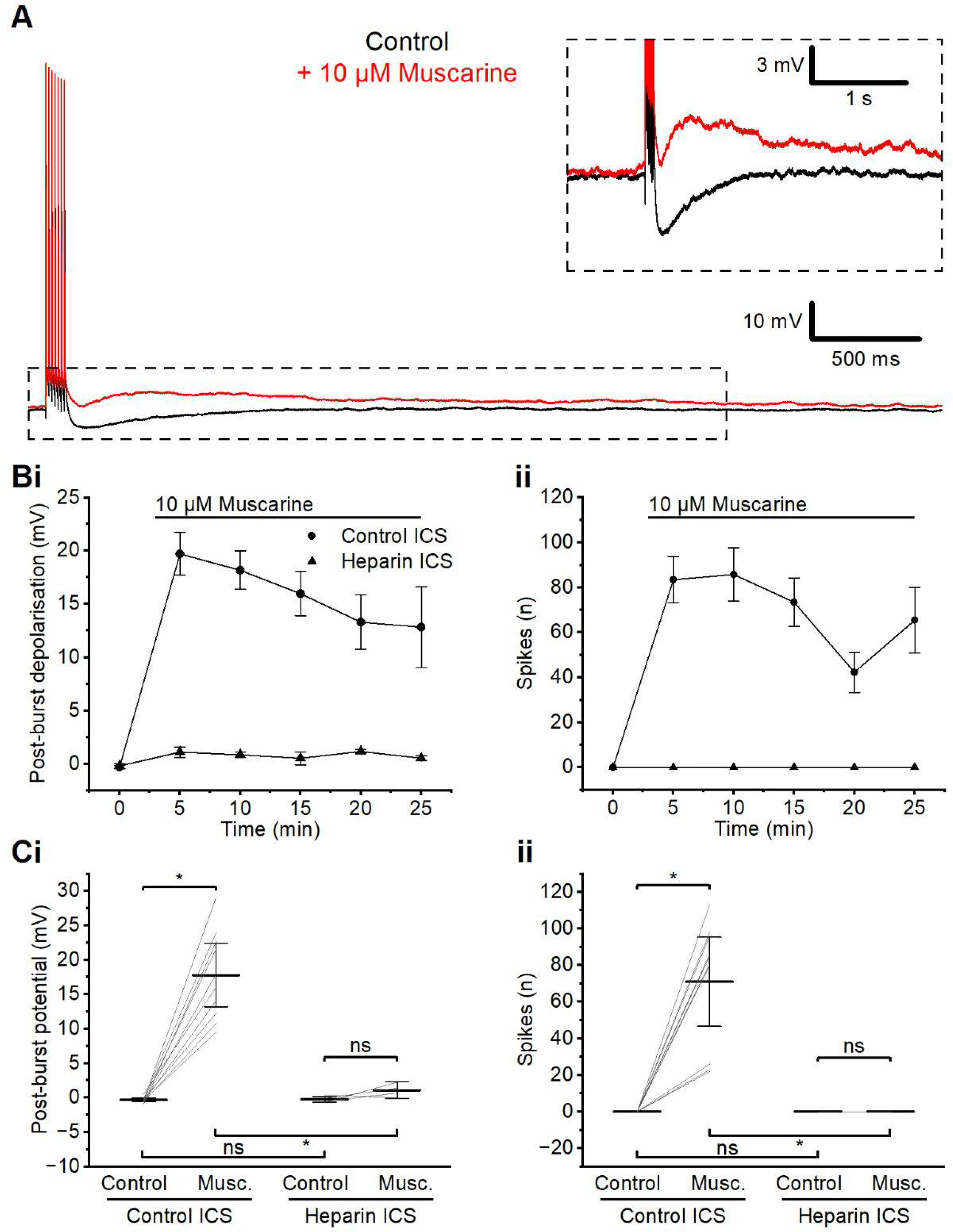
PPs are prevented by a block of IP3 receptors. **A** - Example traces of a cell dialysed with heparin, before and after application of muscarine. The difference in post-bust membrane potential expanded in the inset. **B** - Time course summary of PBD (i) and spikes (ii) following the application of muscarine in cells recorded with control intracellular solution (ICS) and heparin-containing ICS. **C** - Summary plots of the PBD (i) and spikes (ii) before and after the application of muscarine in cells recorded with control ICS and heparin ICS. With control ICS dialysing the cell, wash-in of muscarine significantly changed both PBD and post-burst spiking (WSR test, *n* = 10). When the cell is dialysed with heparin ICS, wash-in of muscarine does not generate spiking PPs, and no significant change in PBD is observed (WSR test, *n* = 4).

**Supplementary figure 5.**
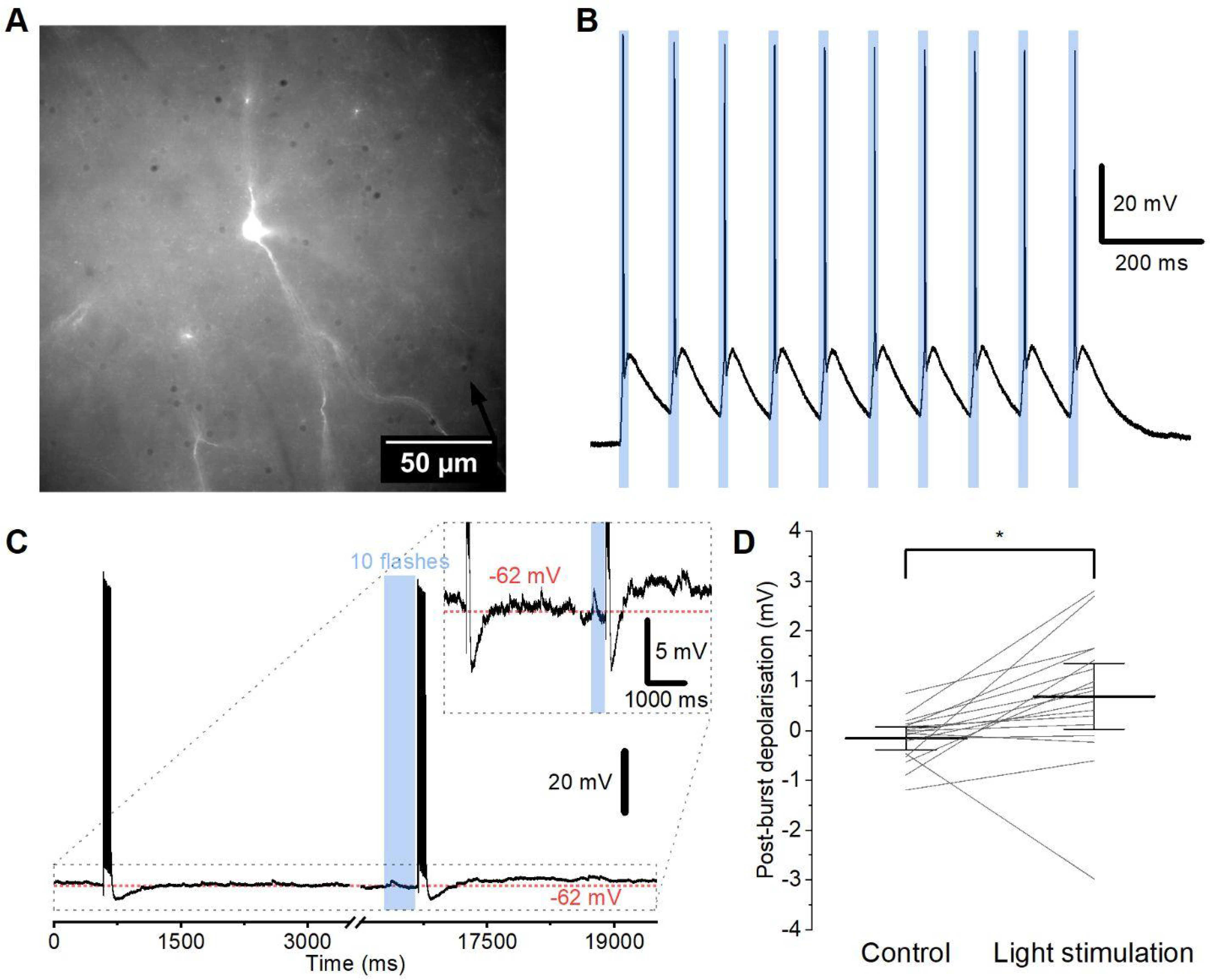
Optogenetic activation of a cholinergic neuron and afferents. **A** - High magnification photomicrograph of an EYFP-positive cell in L2/3 mPFC, with a patch pipette indicated by a black arrow. **B** - Recording from the EYFP-positive (presumably cholinergic) neuron seen in (A), showing the changes in membrane potential during a train of 10 brief flashes (20 ms) of blue light at 25 Hz. **C** - Example traces from an L2/3PC before and after a train of light flashes. The post-burst depolarisation before and after a flash of blue light are shown expanded in the inset. **D** - Summary plot of the PBD amplitude following a train of 10 brief flashes (like in B), presumably causing optogenetic activation of cholinergic afferents. A small, but significant PBD was observed between the control spike-train and spike-train paired with light stimulation (WSR, *n* = 17).

## Statistics

**Figure 2**:

(PBD; 1 µM; Control: -0.100 ± 0.214mV, Muscarine: 0.898 ± 0.254mV; *n* = 10; *p* = 0.0186 by WSR)

(PBD; 3 µM; Control: -0.560 ± 0.288mV, Muscarine: 9.92 ± 2.24mV; *n* = 9; *p* = 0.0039 by WSR)

(PBD; 10 µM; Control: -0.325 ± 0.118mV, Muscarine: 17.8 ± 2.03mV; *n* = 10; *p* = 0.002 by WSR)

(Spikes; 1 µM; Control: 0 ± 0, Muscarine: 0 ± 0; *n* = 10; *p* = NA)

(Spikes; 3 µM; Control: 0 ± 0, Muscarine: 69.3 ± 19.3; *n* = 9; *p* = 0.0312 by WSR)

(Spikes; 10 µM; Control: 0 ± 0, Muscarine: 71.0 ± 10.78; *n* = 10; *p* = 0.002 by WSR)

(PBD; 3 µM vs 10 µM; *p* = 0.0535 by MW) (PBD; 1 µM vs 3 µM; *p* = 0.0016 by MW) (PBD; 1 µM vs 10 µM; *p* < 0.0001 by MW)

(Spikes; 3 µM vs 10 µM; *p* = 0.921 by MW) (Spikes; 1 µM vs 3 µM; *p* = 0.0022 by MW) (Spikes; 1 µM vs 10 µM; *p* < 0.0001 by MW)

**Figure 3**:

(PBD change; Control: -1.77 ± 0.96mV ; *n* = 10)

FFA: -6.24 ± 1.36mV; *n* = 6; *p* = 0.0225 by MWU) ML 204: -2.64 ± 1.4mV; *n* = 8; *p* = 0.829 by MWU)
AC 1903: -5.01 ± 1.3mV; *n* = 6; *p* = 0.118 by MWU)
ML 204 & AC 1903: -13.6 ± 2.59; *n* = 6; *p* = 0.0005 by MWU)

(Spike change; Control: 5.4 ± 9.8; *n* = 10)

FFA: -48.2 ± 15.4; *n* = 6; *p* = 0.01 by MWU)
ML 204: -25.8 ± 10.1; *n* = 8; *p* = 0.036 by MWU)
AC 1903: -51.5 ± 13.9; *n* = 6; *p* = 0.0049 by MWU)
ML 204 & AC 1903: -62.7 ± 11.9; *n* = 6; *p* = 0.0017 by MWU)

**Figure 4**:

(PBD; 2.0 Ca; Control: -0.3252 ± 0.1183, muscarine: 17.8 ± 2.026; *n* = 10; *p* = 0.002 by WSR)

(PBD; 0.1 Ca; Control: -0.5693 ± 0.2045, muscarine: 0.4357 ± 0.3704; *n* = 6; *p* = 0.0313 by WSR)

(PBD; 2.0 vs 0.1: 0.0002 by MWU)

(Spikes; 2.0 Ca; Control: 0 ± 0, muscarine: 71 ± 10.78; *n* = 10; *p* = 0.002 by WSR)

(Spikes; 0.1 Ca; Control: 0 ± 0, muscarine: 0 ± 0; *n* = 6; *p* = NA by WSR)

(Spikes; 2.0 vs 0.1: 0.0002 by MWU)

(PBD; 2.0: 19.58 ± 1.603, 0.1: 7.153 ± 3.349; *n* = 8; *p* = 0.0156 by WSR)

(Spikes; 2.0: 90.1 ± 13.2, 0.1: 26 ± 16.1; *n* = 8; *p* = 0.0078 by WSR)

**Figure 5**:

(PBD; Soma; Pre-puff: -0.22+0.24mV, post-puff: 7.811+2.379mV; *n* = 8; *p* = 0.0078 by WSR) (PBD; Dendrite; Pre-puff: -0.27 ± 0.18mV, post-puff: 2.45 ± 2mV; *n* = 9; *p* = 0.02 by WSR) (Spikes; Soma; Pre-puff: 0+0, post-puff: 26.71+15.65; *n* = 8; *p* = 0.1250 by WSR)

(Spikes; Dendrite; Pre-puff: 0 ± 0, post-puff: 22.1 ± 21.9 ; *n* = 9; *p* = 0.5 by WSR)

PBD; Post-puff soma vs post-puff dendrite; *p* = 0.01828 by MWU

Spikes; Post-puff soma vs post-puff dendrite ; *p* = 0.221 by MWU

(Puff size; Soma; Ca2+: 248,6 ± 16,25 ; *n* = 7

Control: 216 ± 26,94 ; *n* = 6)

(Puff size; Dendrite; Ca2+: 306,7 ± 48,73 ; n = 9

Control: 205 ± 12,58 ; *n* = 4)

**Figure 6**:

(PBD; Cut dendrite; Control: -0.2933 ± 0.07519mV, Muscarine: 7.168 ± 1.581; *n* = 17; *p* = <0.0001 by WSR)

(Spikes; Cut dendrite; Control: 0.05882 ± 0.05882, Muscarine: 59.06 ± 16.3; *n* = 17; *p* = 0.0005 by WSR)

(PBD; Uncut dendrite; Control: -0.253 ± 0.266mV, Muscarine: 13 ± 1.456mV; *n* = 6; *p* = 0.0312 by WSR)

(Spikes; Uncut dendrite; Control: 0 ± 0, Muscarine: 82.83 ± 21.9; *n* = 6; *p* = 0.0312 by WSR)

(Uncut dendrite vs cut dendrite PBD; *p* = 0.0243 by MWU) (Uncut dendrite vs cut dendrite spikes; *p* = 0.1937 by MWU) ( Spearman’s ρ = -0.1833, *n* = 17)

**Suppl. Figure 1:**

(PBD; Control: -0.41 ± 0.1, tACPD: 4.57 ± 1.1; *n* = 12; *p* = 0.0005 by WSR)

(Spikes; Control: 0 ± 0, tACPD: 22.7 ± 7.4; *n* = 12; *p* = 0.0078 by WSR)

**Suppl. Figure 2:**

PBD; soma; control: -5.66 ± 1.52mv, *n* = 6, FFA: -5.91 ± .58mV, *n* = 10; *p* > 0.999 by MWU

PBD; dendrite; control: -2.28 ± 1.4mV, *n* = 3, FFA: -2.12 ± 0.62, *n* = 5; *p* > 0.999 by MWU

Spikes; soma; control: -29.5 ± 8.7, *n* = 6, FFA: -44.9 ± 12.4, *n* = 10; *p* = 0.446 by MWU

Spikes; dendrite; control: -3.67 ± 12, *n* = 3, FFA: -9 ± 5.54, *n* = 5; *p* = 0.929 by MWU

**Suppl. Figure 3:**

PBD change; Control: -1.78 ± 0.96 mV; *n* = 10;

CdCl: -10.9 ± 1.7mV; *n* = 6; *p* = 0.003 by MWU

NiCl: -1.44 ± 1.6mV; *n* = 5; *p* = 0.768 by MWU

Nifedipine: –1.99 ± 0.88mV; *n* = 5; *p* = 0.767 by MWU PD-173212: -0.77 ± 1.1mV; *n* = 7; *p* = 0.42 by MWU

Spike change; Control: 5.4 ± 9.8; *n* = 10;

CdCl: -94.7 ± 12; *n* = 6; *p* = 0.0002 by MWU

NiCl: -29.2 ± 16; *n* = 5; *p* = 0.11 by MWU

Nifedipine: -18.2 ± 8.4; *n* = 5; *p* = 0.35 by MWU

PD-173212: -10 ± 8.9; *n* = 7; *p* = 0.35 by MWU

**Suppl. Figure 4:**

(PBD; control ICS; Control: -0.3252 ± 0.1183 mV, Muscarine: 17.8 ± 2.026 mV, ; *n* = 10; *p* = 0.002 by WSR)

(PBD; heparin ICS; Control: -0.2136 ± 0.2483 mV, Muscarine: 1.088 ± 0.4822 mV, ; *n* = 4; *p* = 0.25 by WSR)

(Spikes; control ICS; Control: 0 ± 0, Muscarine: 71 ± 10.78, ; *n* = 10; *p* = 0.002 by WSR) (Spikes; heparin ICS; Control: 0 ± 0, Muscarine: 0 ± 0, ; *n* = 4; *p* = NA by WSR)

(PBD; control; control ICS vs heparin ICS; *p* > 0.9999 by MWU)

(PBD; muscarine; control ICS vs heparin ICS; *p* = 0.002 by MWU; Hedges’ g = -2.81)

(Spikes; control; control ICS vs heparin ICS; *p* > 0.9999 by MWU)

(Spikes; muscarine; control ICS vs heparin ICS; *p* = 0.002 by MWU; Hedges’ g = -2.25)

**Suppl. Figure 5:**

hPBD; Control: -0.15 ± 0.11mV, Light stimulation: 0.69 ± 0.32mv; *n* = 17; *p* = 0.0046 by WSR

